# Parameter estimation and identifiability analysis of stability and tipping points in potentially bistable ecosystems

**DOI:** 10.64898/2026.03.09.710717

**Authors:** Dasuni Amanda Salpadoru, Matthew P. Adams, Kate Helmstedt, David J. Warne

## Abstract

Ecological regime shifts are potentially a common property of ecosystems, describing transitions between alternative stable states that can represent healthy or unhealthy conditions under the same environmental drivers. Once a tipping point, defined as a critical threshold separating alternative stable states, is crossed, the system may degrade and recovery can be difficult, making early detection essential for effective ecosystem management. Predicting these tipping points requires models that exhibit bistability, representing systems that can exist in two alternative stable states under identical environmental conditions. A key question is whether standard ecological monitoring data can be used to identify bistability and accurately estimate tipping points. Using the Carpenter model of lake eutrophication, which expresses bistability between clear and polluted water states, we generate synthetic data under known stability regimes. Profile likelihood analysis is then applied to assess parameter identifiability and detect system stability and tipping points. Our results show that standard monitoring data do not always provide sufficient information to distinguish bistable from stable regimes. Importantly, bistability and tipping points become practically identifiable only when data are collected very close to the tipping point.

## 1. Introduction

Ecosystems are complex and dynamic systems that respond to gradual environmental changes, such as human activities, rising temperatures, and increasing nutrient loading, in different ways. These responses may be linear and continuous, with state variables changing smoothly as conditions vary (Figure 1(a)). In contrast, nonlinear dynamics can generate alternative stable states under identical environmental conditions, with outcomes depending on initial conditions and nonlinear interactions (Figure 1(b)).

**Figure 1.**
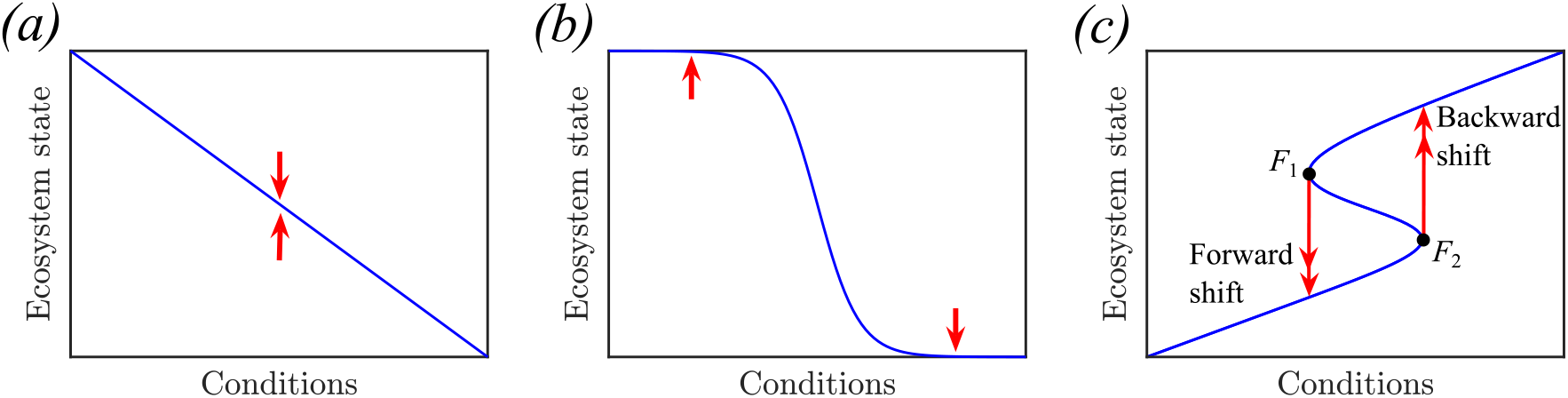
Ecosystem state at equilibrium can be affect by environmental conditions in different ways. In response to changes in environmental conditions the system at equilibrium can change: (a) linearly; (b) nonlinearly with substantial, but reversible, change occurring within a small range of environmental changes; and (c) nonlinearly and irreversibly due to the effect of bistability. In this bistable case the system exhibits multiple equilibria withing a range fo environmental conditions (between *F*_1_ and *F*_2_). If the system in (c) is on the upper branch, close to the *F*_1_ bifurcation point, a small gradual change can push the system past a threshold, triggering a sudden transition to the lower stable state (forward shift). A backward shift happens if conditions are reversed sufficiently to cross the second bifurcation point, *F*_2_.

Nonlinear interactions can lead the system to exhibit more than one stable state. When exactly two stable states coexist, separated by an unstable equilibrium, small perturbations or environmental drivers can cause the system to shift suddenly from one state to the other. The system can then stabilise in either of the two states and transition suddenly when thresholds are crossed. This type of behaviour is known as bistability (Figure 1(c)) (Scheffer et al., 2001). In bistable systems, the unstable equilibrium represents a tipping point, defined as the critical threshold at which small, gradual changes accumulate to result in a large and rapid change in system state. Crossing a tipping point, either through gradual environmental change or sudden perturbations, results in a regime shift, defined as the transition of the ecosystem from one stable state to another (Scheffer and Carpenter, 2003). Such regime shifts are often difficult to reverse because the system response to changes in external conditions differently depending on its past trajectory. The property, known as *hysteresis*, causes a great challenge in restoring ecosystems as reversing environmental conditions to their previous levels need not return the system to its original state.

Accurate prediction of tipping points and regime shifts not only depends on understanding the ecosystem dynamics, but also on reliable estimation of model parameters. This is critical because tipping point estimates depend on accurate parameter values, and if the available monitoring data do not provide sufficient information to estimate these parameters accurately, predictions of critical thresholds and system stability can be misleading. Such missclassification of stability has serious implications in bistable settings: once a tipping point is crossed, returning to the previous state may be difficult, even if environmental conditions are restored (Wu et al., 2019).

Bistability has been documented across a wide range of ecosystems, including lakes that can alternate between clear states dominated by submerged vegetation and turbid states dominated by phytoplankton, coral reefs that shift from coral-dominated communities to systems dominated by fleshy macroalgae, woodlands that transition between herbaceous vegetation and woody plant dominance, and desert ecosystems that alternate between perennial vegetation and bare soil with ephemeral plants (Scheffer et al., 2001; Carpenter et al., 1999). These gradual environmental changes can trigger sudden regime shifts with significant ecological consequences (Scheffer et al., 2001; Scheffer and Carpenter, 2003; Wu et al., 2019). Beyond ecology, bistability is an essential concept in the study of biochemical systems (Jaruszewicz-Błońska and Lipniacki, 2017; Lugagne et al., 2017; Tian and Burrage, 2006; Warne et al., 2019), engineering (Barakat et al., 2025; Cao et al., 2021; Podder et al., 2016) and particle physics (Lee, 2018; Tolle et al., 2025; Zhang et al., 2025).

Among these examples, we focus on lake ecosystems because freshwater ecosystems are increasingly threatened by eutrophication, a critical environmental problem affecting over 40% of lakes globally due to increased phosphorus levels (Vinçon-Leite and Casenave, 2019). Eutrophication is recognized as one of the primary threats to freshwater quality (Smith et al., 2006). Lake eutrophication is a process caused by excessively high concentrations of nutrients, primarily phosphorus and nitrogen, within the water body (Akinnawo, 2023). The source of these excess nutrients primarily originate from human activities, including agricultural runoff, untreated sewage discharge, and industrial waste (Devlin and Brodie, 2023). Although natural eutrophication can occur gradually over centuries (Lin et al., 2021), the accelerated form driven by human activity, anthropogenic eutrophication, is much more rapid (Smith and Schindler, 2009). One of the most visible consequences of eutrophication is increased phytoplankton biomass, which reduces light penetration and can lead to the loss of benthic vegetation. Light limitation can harm aquatic organisms, affect nutrient cycling, biodiversity, and water quality (Schindler et al., 2008; Hart et al., 2025; Bhagowati and Ahamad, 2019; Lin et al., 2021). Due to these broad ecological impacts, eutrophication has become a central topic in freshwater ecosystem research.

Mathematical models provide a well characterized framework for exploring the phosphorus cycle in lake ecosystems (Carpenter et al., 1999). Nutrient enrichment can cause a sudden transition from a low phosphorus level (oligotrophic) to a high phosphorus level (eutrophic) once a critical threshold in phosphorus concentration is crossed (Scheffer et al., 2001). In this context, the oligotrophic and eutrophic conditions correspond to the two alternative ecological regimes, each representing a stable long-term state of the system. This behavior illustrates bistability, where the lake can exist in either the oligotrophic or eutrophic state under the same environmental conditions (Robertson et al., 2003; Bhagowati and Ahamad, 2019). In such systems, equilibrium points correspond to stable steady states where phosphorus concentrations remain relatively constant over time. To investigate these dynamics, theoretical and modelling studies have provided understanding into the dynamics that drive regime shifts in bistable ecosystems (Scheffer et al., 1993; Wu et al., 2019; Carpenter et al., 1999; Boada et al., 2017).

While long-term monitoring can indicate whether a lake is experiencing eutrophication or maintaining a clear-water state, it does not reveal whether the system dynamics correspond to a bistable or stable region (Stow et al., 1998; Wu et al., 2019; Ausseil et al., 2024; Canfield Jr. et al., 2002). This is important in bistable systems, where two alternative stable states can exist under the same environmental conditions, and hysteresis can occur—meaning the system’s response depends on its past trajectory, with forward and backward transitions occurring at distinct thresholds (Carpenter et al., 1999; Scheffer et al., 1993). To detect such transitions, monitoring data must contain sufficient information to estimate the model parameters that drive these shifts. This limitation is known as parameter identifiability, which refers to the ability to estimate model parameters uniquely from observed data (Browning et al., 2020; Raue et al., 2009; Gutenkunst et al., 2007). Formally, a model is identifiable when different parameter values lead to distinct observable outcomes; with ideal data, the true values of all parameters could be uniquely determined (Hines et al., 2014; Raue et al., 2009; Roosa and Chowell, 2019). Parameter identifiability is therefore essential for reliable inference: without it, even extensive monitoring data may not allow accurate estimation of parameters or prediction of critical transitions. Poor identifiability increases uncertainty in predicted tipping points and can lead to misclassification of system stability, reducing confidence in model-based predictions (Moore, 2018; Carpenter and Brock, 2006). Understanding which parameters most strongly influence system behaviour is therefore important for reliable predictions of regime shifts and for guiding effective ecosystem management (Scheffer et al., 2001; Wu et al., 2019).

In this study, we address the question of whether bistability and tipping points can be identified from practically measurable data. We generate synthetic datasets representing both bistable and non bistable region which represents a system with a single stable equilibrium state under varying levels of observational noise, using the Carpenter model of lake eutrophication (Carpenter et al., 1999). While it is well established that lakes and other ecosystems can exhibit bistability and tipping points, the ecological modelling literature has not yet clarified whether typical monitoring data are sufficient to detect bistability or to reliably estimate the parameters that define these dynamics (Wu et al., 2019).

To address this challenge, we apply profile likelihood analysis (Section 2.4) (Raue et al., 2009; Pawitan, 2001; Simpson et al., 2022) that follows the principles of likelihood-based frequentist inference to assess parameter identifiability, combined with a novel statistical framework (Section 2.5) designed to evaluate the stability classification of each model parameter and determine whether it can reliably indicate bistable or stable behavior in lake ecosystems. Using these approaches, we evaluate parameter identifiability and quantify how uncertainty in model parameters influences our ability to estimate system stability. Our results provide concrete guidance for ecological monitoring by demonstrating that reliable detection of regime shifts requires data collected close to tipping points, where system sensitivity to parameter variation is highest. Conversely, monitoring strategies that sample too far from these critical thresholds may fail to identify bistability, even when model parameters are theoretically identifiable. Overall, this work presents and demonstrates a general methodology for identifying the potential presence of alternative stable states and tipping points in ecosystems for which observational data and a mechanistic model are available.

## 2. Materials and Methods

In this paper, we aim to investigate the parameter identifiability of the bistable dynamics in lake ecosystems using realistic monitoring data (Section 2.1). To achieve this, first we analyse parameter identifiability within ecological models that are capable of exhibiting bistable behavior, focusing on the lake eutrophication model (Section 2.2). This model describes the phosphorus cycle process in lakes, which contain stable or bistable behaviour depending on the values of model parameters. Our analysis focuses on assessing the identifiability of model parameters across stable and bistable regions, under different equilibrium states and varying levels of observational noise, as well as on determining the identifiability of ecosystem stability from the corresponding synthetic datasets. We use a standard statistical tool, maximum likelihood estimation, to estimate model parameters. To address parameter identifiability and associated uncertainty, we apply profile likelihood-based identifiability analysis (Section 2.4).

### 2.1. Lake monitoring data

Long-term lake monitoring data are used to identify patterns in lake behavior and understand changes in actual phosphorus concentration over time (Lathrop et al., 1998). Regular monitoring is important for detecting long-term trends in nutrient dynamics and informing management strategies aimed at preventing eutrophication (Stow et al., 1998). In real-world applications, sampling frequencies vary considerably depending on the lake’s purpose and monitoring resources. Typically, monitoring focuses on the biologically active season, with chlorophyll a measurements taken several times per year as an indicator of phytoplankton biomass. In contrast, other biological components such as aquatic plants collect less frequently, often only once per year (Lyche-Solheim et al., 2013). For instance, Lake Mendota has been monitored annually to study long-term phosphorus trends (Lathrop et al., 1998), while Florida lakes are monitored monthly for total phosphorus, total nitrogen, and chlorophyll-a concentrations (Canfield Jr. et al., 2002).

Figure 2 illustrates a simulated time series showing how phosphorus levels change under these assumptions, displaying a characteristic sigmoid pattern. In this study, we explore questions of parameter and stability identifiability using simulated noisy data inspired by typical lake monitoring programs. Following the approach of Guttal and Jayaprakash (2007), we simulate biannual phosphorus measurements based on long-term monitoring practices in lakes such as Lake Mendota (Lathrop et al., 1998). Details of specific simulated data scenarios we consider is described in Section 2.6.

**Figure 2.**
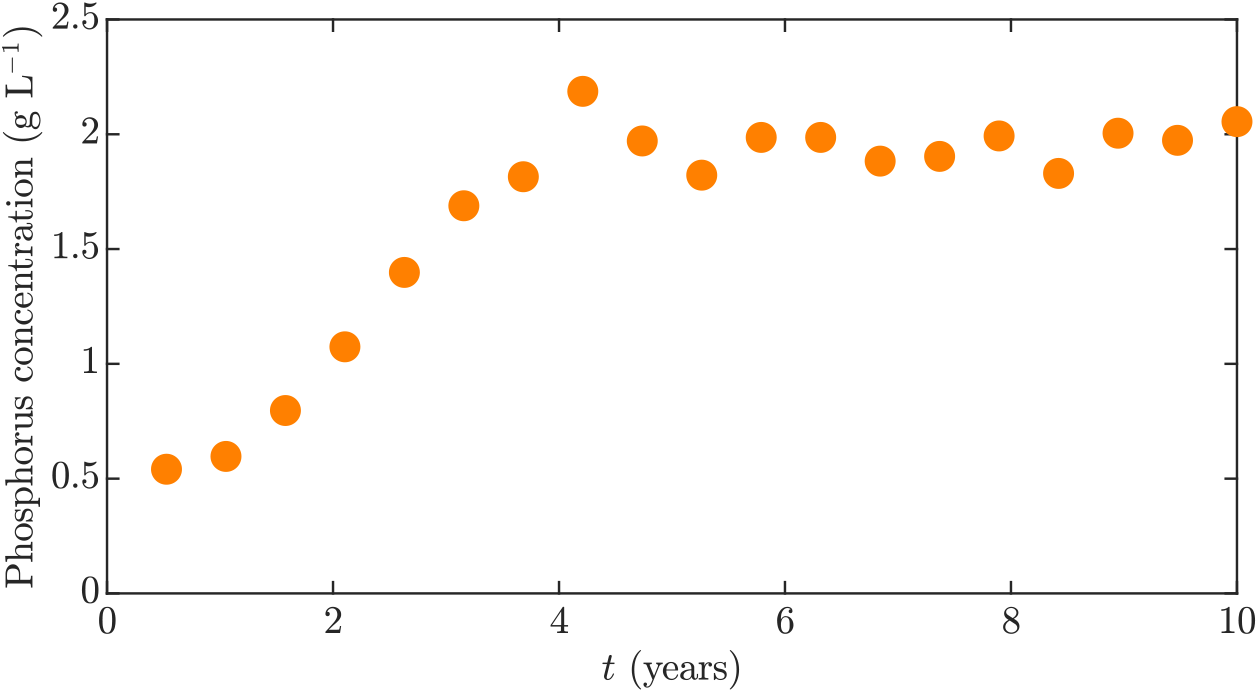
Simulated time series representing typical data obtained from lake monitoring programs, showing noisy measurements of phosphorus concentration (orange circles) on a biannual basis. The times series is simulated using the Carpenter model of lake eutrophication (Carpenter et al., 1999) that is presented in Section 2.2. The simulation depicts the eutrophication of a lake that is characterised by a sigmoidal increase in phosphorus toward the eutrophic stable equilibrium that occurs at a phosphorus concentration of approximately 2 (g L^−1^)

### 2.2. Mathematical modelling

We consider the lake eutrophication model developed by Carpenter et al. (1999), which describes the phosphorus cycle process in freshwater lakes. The model was driven by three core processes: *(i)* phosphorus loading, *(ii)* phosphorus sinks, and *(iii)* phosphorus recycling. Phosphorus loading refers to the input of phosphorous into the lake primarily through external loading from the surrounding watershed, such as agricultural run-off or urban discharge. Phosphorus sinks refer to the portion of the phosphorus that is removed through physical processes including sedimentation or outflow. Phosphorus recycling refers to the accumulation of phosphorus that is not permanently lost, but reintroduced through processes such as resuspension of sediments, chemical processes, and microbial absorption. These recycling process can lead to internal phosphorus loading that can compound the amount of phosphorus in the water column (Carpenter et al., 1999; Genkai-Kato and Carpenter, 2005; Houser et al., 2000; Rychła et al., 2014). These recycling mechanisms can lead to a positive feedback loop that generates alternative stable states in lake ecosystems. The balance among these processes determines the overall phosphorus status of the lake ecosystem (Carpenter and Brock, 2006).

This model has been widely used to study phosphorus dynamics in freshwater ecosystems (Carpenter et al., 1999), and it is a canonical model example for studying regime shifts (Guttal and Jayaprakash, 2007; Wu et al., 2019). The model is phenomenological rather than fully mechanistic, and describes the phosphorus cycle using a small number of parameters that summarise complex biogeochemical processes. This makes it computationally efficient for parameter estimation and identifiability analysis. These key dynamics of lake eutrophication can be described using the Carpenter model (Carpenter et al., 1999) that consists of a non-linear ordinary differential equation (ODE),

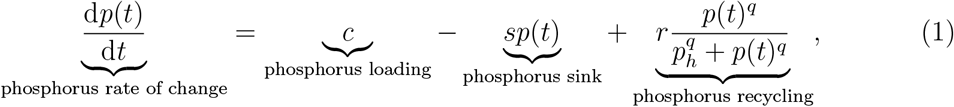

where *p*(*t*) ≥ 0 (g L^−1^) is the phosphorus concentration at time *t* ≥ *t*_0_ (years), with initial time *t*_0_ ≥ 0. The phosphorus loading rate is *c* ≥ 0 (g L^−1^ year^−1^), the phosphorus sink rate is *s* ≥ 0 (year^−1^), the maximum internal recycling rate is *r* ≥ 0 (g L^−1^ year^−1^), the nondimensional Hill coefficient is *q* ≥ 2, which controls the steepness of the phosphorus recycling response curve, and *p*_*h*_ ≥ 0 (g L^−1^) is the phosphorus concentration at which the recycling rate become half maximum. The system is initialized with the condition *p*(*t*_0_) = *p*_0_, where *p*_0_ ≥ 0 is the phosphorus concentration at the starting time *t*_0_. A summary of all model parameters and variables is provided in Table 1.

**Table 1:** Description of model parameters for the Carpenter model of lake eutrophication.

| Parameter/variable | Description | Units |
| --- | --- | --- |
| $p(t)$ | Phosphorus concentration in the water column | $\text{g L}^{-1}$ |
| $c$ | External phosphorus loading rate | $\text{g L}^{-1} \text{year}^{-1}$ |
| $s$ | Phosphorus sink rate | $\text{year}^{-1}$ |
| $r$ | Maximum internal phosphorus recycling rate | $\text{g L}^{-1} \text{year}^{-1}$ |
| $p_h$ | Phosphorus concentration at half-maximum recycling | $\text{g L}^{-1}$ |
| $q$ | Hill coefficient | — |
| $t$ | Time | year |

An important feature of the model (Equation (1)) is the behaviour of the nonlinear Hill term, which captures how phosphorus recycling responds to changes in phosphorus concentration. The Hill coefficient *q* controls the sensitivity of recycling to phosphorus levels, with *p*_*h*_ representing the concentration at which recycling reaches half of its maximum. At low phosphorus levels, recycling remains minimal; however, beyond a tipping point, the phosphorus concentration increases rapidly (Appendix A). This nonlinear response gives rise to two distinct stable states, highlighting a characteristic of bistable dynamics (Guttal and Jayaprakash, 2007).

One can obtain some initial intuition on how the stability of the Carpenter model (Equation (1)) is affected by parameters using the bifurcation diagram provided in Figure 3. Following Guttal and Jayaprakash (2007), this diagram is constructed by numerically solving for *p*_∞_ such that,

**Figure 3.**
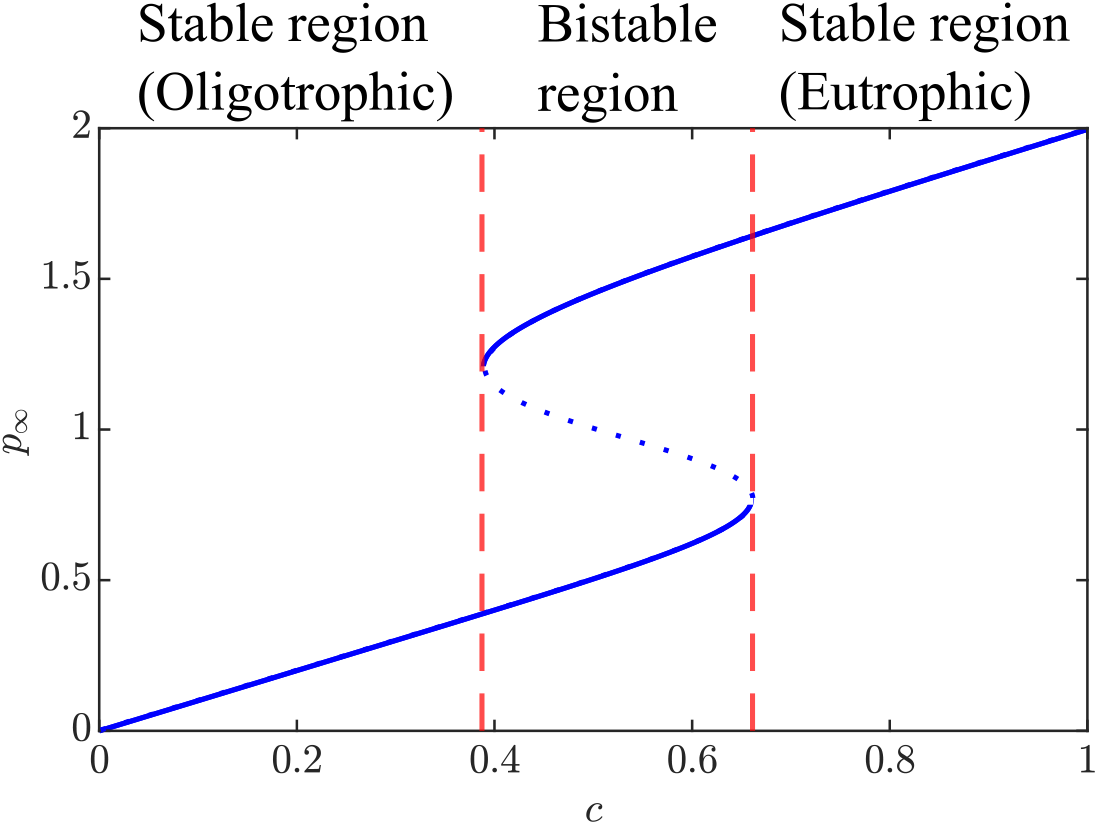
The bifurcation diagram for the Carpenter model (Equation (1)) shows the potential steady state phosphorus concentration *p*_∞_, given by solutions to Equation (2) (blue line), along with is dependence on the phosphorus loading rate *c*. The curve shows regions of stability and bistability with *c* values indicating regime shifts indicated (dash red lines at *c* = 0.39 and *c* = 0.66). In the bistable regime, the system is attracted to one of the two alternative stable equilibria depending on the whether the initial phosphorus concentration is above or below the tipping point (blue dotted line). The parameters are fixed as *s* = 1 (year^−1^), *r* = 1 (year^−1^), *q* = 8, *p*_*h*_ = 1 (g L^−1^), with varying *c* ∈ [0, 1] (g L^−1^ year^−1^).

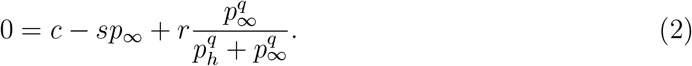

For fixed values of the sink rate *s* = 1 (year^−1^), maximum recycling rate, *r* = 1 (year^−1^), and Hill function variables, *p*_*h*_ = 1 (gL^−1^) and *q* = 8, we see their are three distinct stability regimes depending on the value of the loading rate *c* ∈ [0, 1] (gL^−1^year^−1^). For *c* < 0.39 (gL^−1^year^−1^) we have an unconditionally stable systems that approaches a healthy oligotrophic state. In contrast for *c* > 0.66 (gL^−1^year^−1^) the system remains unconditionally stable, however, it approaches the unhealthy eutrophic state. Between these two stable regions, for 0.39 (gL^−1^year^−1^) < *c* < 0.66 (gL^−1^year^−1^), we have a bistable regime in which there are two stable equilibria (the lower representing oligotrophic and the upper representing eutrophic) that are separated by an unstable equilibrium, representing a tipping point (Scheffer et al., 2001; Scheffer and Carpenter, 2003). If the phosphorus concentration is perturbed such that it crosses the tipping point, then the stable equilibrium the system evolves toward will switch. Of course the specific thresholds for the loading, *c*, will depend on the specific values of *s, r, p*_*h*_ and *q* (See Appendix A). As a result, determining the stability regime and potential tipping points requires parameter estimation and uncertainty quantification.

### 2.3. Parameter estimation

The exploration of the bifurcation diagram (Figure 3 and Appendix A) highlights that parameter values play a critical role in determining whether a lake ecosystem is bistable or stable. To understand and predict these dynamics, especially near tipping points, it is necessary to estimate the model parameters accurately. Therefore, estimating the parameter set is important for linking the mathematical model to real-world phosphorus data and for identifying where regime shift occur (Figure 3).

Statistical inference in dynamic ecological models commonly uses both Bayesian (Wu et al., 2019; Hines et al., 2014) and frequentist (Simpson et al., 2022; Raue et al., 2009) frameworks for parameter estimation and uncertainty quantification. In this study, we use a frequentist approach, applying maximum likelihood estimation (MLE) to estimate model parameters. Bayesian methods can be computationally intensive for prescribing prior distributions, and are generally more advantageous for models with higher dimensional parameter spaces. Since our model has a relatively small number of parameters, MLE is unlikely to become trapped in local likelihood maxima, making the frequentist approach both efficient and appropriate.

We denote *y*_*i*_ as the phosphorus concentration observed at discrete times *t*_*i*_, where *i* = 1, 2, …, *n* and *n* is the total number of observations. These measurements are subject to observational noise and are distinguished from the model-predicted values, *p*_*i*_ = *p*(*t*_*i*_; ***θ***), where ***θ*** = (*c, s, r, q, p*_*h*_, *σ*) represents the parameters of the Carpenter model (Equation (1)) that are to be estimated including the observation standard deviation *σ* > 0. To estimate model parameters (***θ***), we assume that the observations are noisy versions of the model predictions and that they are normally distributed,

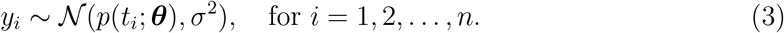

We assume homoskedastic additive Gaussian noise that is conditionally independent of the ODE solution *p*(*t*_*i*_; ***θ***). Each observation deviates from the model prediction according to a normal distribution with variance *σ*^2^, representing the magnitude of measurement error and other small-scale variability not captured by the model. Because this variance reflects the overall uncertainty in the data, it is estimated alongside the other model parameters and is therefore included in the parameter vector ***θ***.

To estimate the parameters ***θ***, the standard frequentist approach uses the maximum likelihood test. Let 𝒟= { (*t*_1_, *y*_1_), (*t*_2_, *y*_2_), …, (*t*_*n*_, *y*_*n*_) } denote the dataset of observations *y*_*i*_ recorded at times *t*_*i*_. The likelihood function which is the probability density function for observing the data 𝒟 under the assumption of our model (Equation (1) and Equation (3)) with parameters ***θ***, is defined as

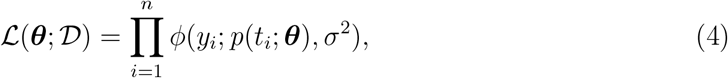

where *ϕ*(*x*; *µ, σ*^2^) denotes the Gaussian probability density function with mean *µ* and variance *σ*^2^. To simplify differentiation calculations and improve numerical stability, we define the log-likelihood function, that is,

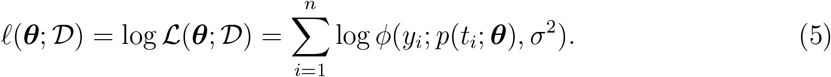

To estimate the parameters ***θ*** using the MLE, we identify the parameter values that maximise the log-likelihood function. Formally, the MLE is defined as

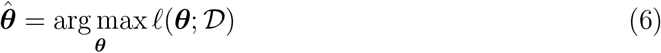

If 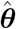 is the MLE of the model parameters ***θ*** = (*c, s, r, q, p*_0_, *σ*), then, by the invariance property, the MLE of phosphorus trajectory is 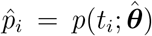 at time *t*_*i*_. The estimation process determines the optimal values for all six parameters, *c, s, r, q, p*_*h*_, and *σ* such that the model predictions closely align with the observed measurements as indicated in Figure 4.

**Figure 4.**
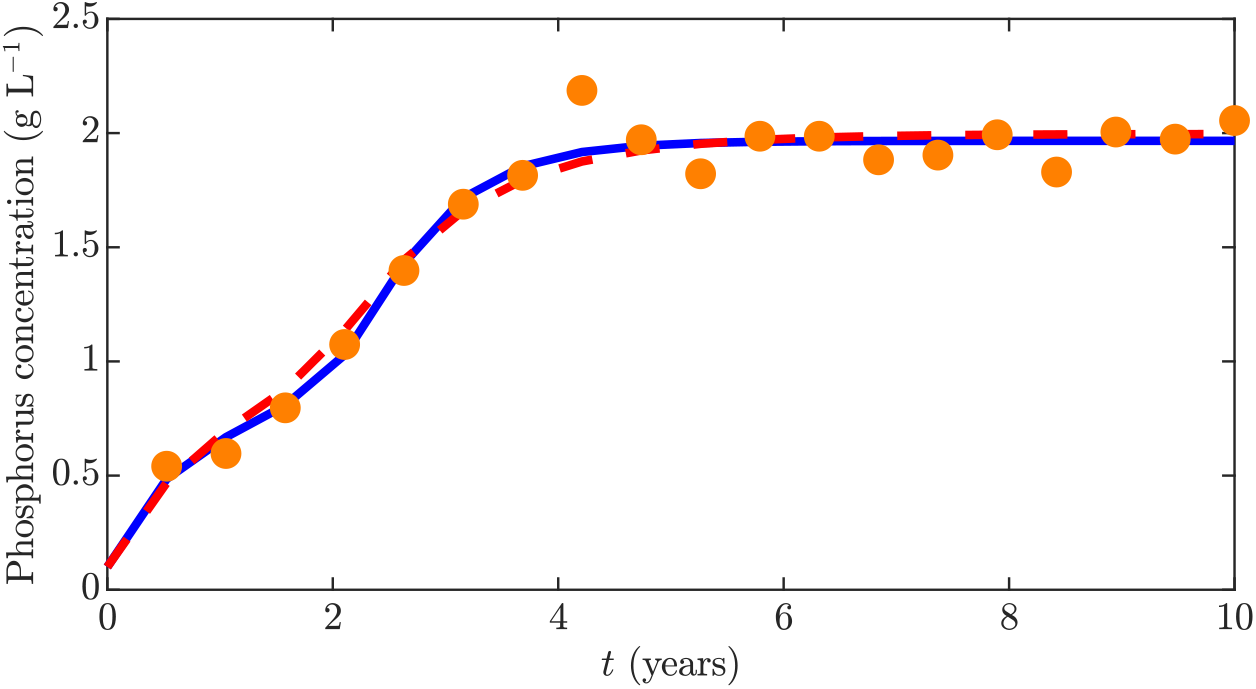
Simulated noisy phosphorus concentration observation (orange circles) together with the underlying true phosphorus trajectory (blue solid line) and the MLE of the phosphorus trajectory (red dashed line; Equation (1)). The MLE trajectory represents the best-fit solution obtained by estimating the parameter vector ***θ*** using maximum likelihood. In this experiment, the true parameter values were *c* = 1 (g L^−1^ year^−1^), *s* = 1 (year^−1^), *r* = 1 (g L^−1^ year^−1^), *q* = 8, *p*_*h*_ = 1 (g L^−1^), and *σ* = 0.1 (g L^−1^). The corresponding maximum likelihood estimates were *c* = 1.2544 (g L^−1^ year^−1^), *s* = 1.6434 (year^−1^), *r* = 2.0000 (g L^−1^ year^−1^), *q* = 6.6914, *p*_*h*_ = 1.0113 (g L^−1^), and *σ* = 0.0892 (g L^−1^).

The parameter estimation process is subject to bound constraints to ensure biologically meaningful values, with parameters restricted such that all parameters are non-negative, that is ***θ*** ∈ [0, ∞)^6^. Estimating 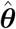 requires maximizing a multidimensional function, which can be achieved through various numerical optimization techniques. In this study, we utilize MATLAB’s `fmincon`, which implements a gradient-based constrained optimization algorithm, allowing the specification of bound constraints and supporting both local and global searches. The local search is initialized with a reasonable estimate of 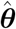, while the global search, computationally more expensive, helps reduce the risk of converging to a local maximum.

### 2.4. Parameter identifiability analysis

Identifiability analysis consists of two key aspects: structural and practical identifiability. Structural identifiability assesses whether a parameter can be uniquely determined based on the model formulation alone, whereas practical identifiability considers the limitations applied by real-world data such as noise and measurement constraints (Pawitan, 2001). Prior to conducting practical identifiability, it is essential to verify structural identifiability of the model. In this work we perform Structural Identifiability analysis using the GenSSI (Generating Series for testing Structural Identifiability) software (Chiş et al., 2011), which determines whether the model structure enables unique parameter estimation in the absence of noise. GenSSI is a MATLAB-based toolbox designed to analyze the structural identifiability of nonlinear dynamic models and is very easy to use (Chiş et al., 2011). It uses generating series and identifiability analysis to automate symbolic computations, helping users determine whether model parameters are structurally globally, locally, or non-identifiable.

A parameter is non-identifiable if it cannot be uniquely determined from the model or data. If a model is found to be structurally non-identifiable, then further practical identifiability analysis is not meaningful, because structural non-identifiability implies that the parameters cannot be uniquely estimated even with ideal, noise-free data, and therefore they cannot be practically identifiable from finite, noisy observations (Guillaume et al., 2019). Structural non-identifiability arises from overparameterization of the model including its model structure or observation function, meaning that parameters cannot be uniquely estimated even with perfect data. In contrast, practical non-identifiability occurs when the model is structurally identifiable but the available data are insufficient, due to noise or limited observations, to reliably estimate the parameters. (Chiş et al., 2011).

To proceed with the practical identifiability analysis, we use the frequentist profile likelihood approach (Raue et al., 2009; Simpson et al., 2022). It is a powerful method for practical identifiability analysis because it is effective with small sample sizes, handles asymmetric confidence sets, and remains stable even in cases of practical nonidentifiability (Pawitan, 2001; Warne et al., 2024).

As described by Simpson et al. (2022), the practical identifiability analysis proceeds by constructing profile likelihoods for each model parameter. The profile likelihood analysis is based on the likelihood function (Equation (4)) and fixes one parameter at a time while optimizing over all others. This provides detailed understanding of how the likelihood varies with respect to the parameter of interest. We assume that the full parameter set ***θ*** can be partitioned into two subsets: the parameter of interest, ***ψ***, and the nuisance parameters, ***η***. That is, ***θ*** = (***ψ, η***). Given a set of data 𝒟, the profile log-likelihood for the parameter of interest ***ψ*** is defined as

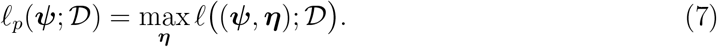

The profile-likelihood approach optimizes out the nuisance parameters ***η*** for a fixed value of the parameter of interest ***ψ***. Once the profile log likelihoods are calculated, they are normalized by subtracting the maximum log-likelihood value,

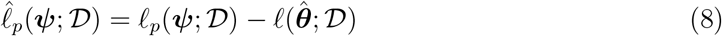

where 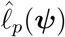 represents the normalized log-likelihood, expressing all profile log likelihood values relative to this maximum. This normalization ensures that likelihood profiles can be directly compared across different parameter values. By the shape of the normalised profile likelihood curves of the model parameters, we can qualitatively decide whether that respective parameter is identifiable or not by look at its curvature (Figure 5). Identifiable parameters are charactised by curves with a distinct maximum (Figure 5(a)), whereas non-identifiable (Figure 5(b)) or partially identifiable (Figure 5(c)) parameters will exhibit flat regions.

**Figure 5.**
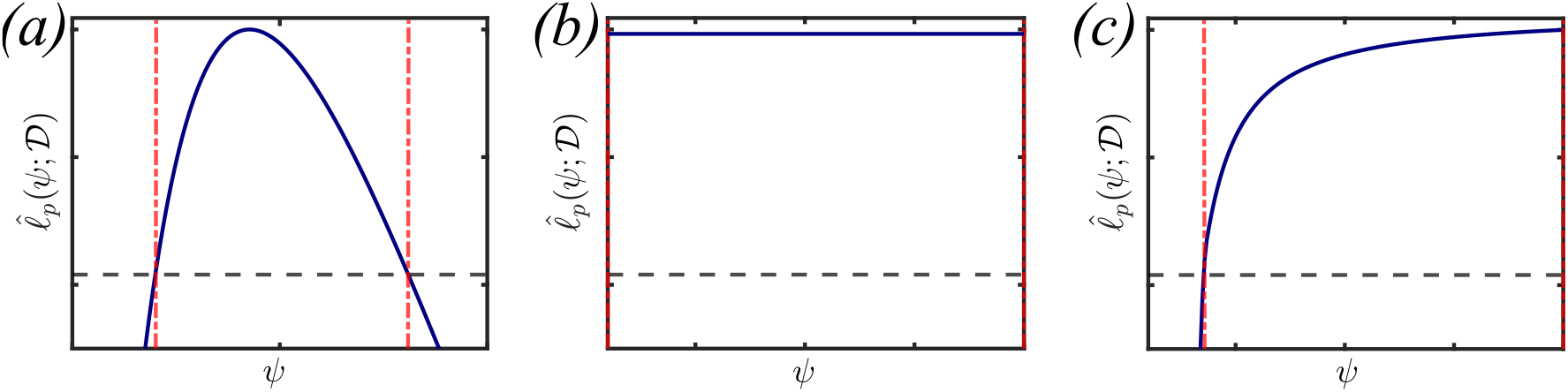
Illustration of the three main types of normalized log-profile likelihood curves (blue curves), along with the upper and lower bounds of the profile-based confidence interval (dashed red lines) and the confidence threshold (dashed black line). (a) Shows a narrow profile curve within a well-defined confidence interval, confirming that the corresponding parameter is identifiable. This implies that the data provide a unique and well-constrained estimate of that parameter. (b) Shows a flat profile curve, indicating nonidentifiability; the data lack sufficient information to estimate the parameter. (c) Displays a partially identifiable curve, meaning the parameter is only identifiable within a limited range, and uncertainty remains in its estimation.

Likelihood profiles can also be used to perform the uncertainty quantification for each parameter, we obtain confidence sets from the normalized profile log-likelihoods. By Wilks’ theorem (Appendix B.2), the likelihood ratio statistic is asymptotically *χ*^2^-distributed (Wilks, 1938; Pawitan, 2001). As a consequence, the confidence region for a parameter ***ψ*** consists of all values satisfying

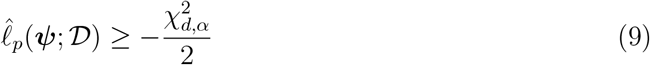

Here, 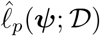 is the profile log-likelihood at ***ψ***, *d* is the number of constrained parameters (that is, the dimensionality of the interest paremeter ***ψ***), and 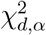 is the upper *α*-quantile of the chi-squared distribution with *d* degrees of freedom. For a 95% confidence level (*α* = 0.05), the critical threshold identifies points that satisfy this criterion and thus contained within the confidence set of the interest parameter. After computing the profile likelihood and confidence set for each parameter, we can confirm the precision with which model parameters can be estimated based on the available data. In addition, we can visualise the resulting profile likelihoods for each parameter.

For the purposes of identifiability analysis in relation to the Carpenter model (Equation 1), we compute univariate profiles using uniform grids of *n* =1,000 points across realistic ranges. Based on the work of Guttal and Jayaprakash (2007), we determine the following ranges to be sensible: *c* ∈ [0, 1] g L^−1^ year^−1^, *s* ∈ [0.1, 2] year^−1^, *r* ∈ [0.1, 2] g L^−1^ year^−1^, *q* ∈ [2, 16], *p*_*h*_ ∈ [0.1, 2] g L^−1^, and *σ* ∈ [0, 0.15] g L^−1^. Note, that more generally there are methods for constructing profile boundaries without prescribing a fixed grid of points (Simpson and Maclaren, 2023), but in our application these are not necessary.

### 2.5. Profile-wise analysis of stability and tipping-points

The likelihood-based approach to parameter identifiability and uncertainty quantification can also be utilised for uncertainty quantification quantities derived from parameters. Following Simpson and Maclaren (2023), the invariance property of the MLE can be exploited to obtain profile-based confidence sets, which can be viewed as a from of sensitivity analysis (Liu et al., 2026). To summarise the general concept, let 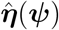 be the values of the nuisance that satisfy Equation (7) to obtain 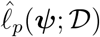. That is,

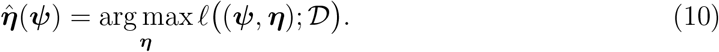

Note that 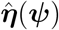 is the MLE for the nuisance parameters for the fixed value of the interest parameter. As a result, for ***ψ*** within the confidence set (statisfying Equation (9)) then any function 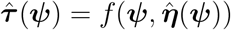 will also be within this profile-based confidence set.

In the setting where 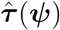 represents a model prediction, Simpson and Maclaren (2023) demonstrate that approximate frequentist prediction sets can be obtained through a union of univariate profile-based confidence sets. In this work, we utilise the profile-wise analysis approach to perform a sensitivity analysis on model stability and tipping points, both of which can be viewed as special cases of 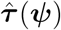.

Given a set of parameters from the Carpenter model ***θ*** = (*c, s, r, q, p*_*h*_, *σ*), we can obtain steady state solutions through setting d*p/*d*t* = 0 and hence solving Equation (2). This is equivalent to finding the real roots of the function,

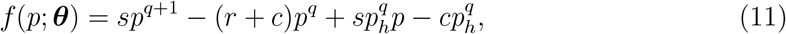

so that the set of steady states, for a given parameter set, *P* (***θ***) = {*p* : *f* (*p*; ***θ***) = 0, *p* ∈ [0, ∞)}. Thus *p*_∞_ is a steady state if and only if *p*_∞_ ∈ *P* (***θ***).

For each steady state, *p*_∞_ ∈ *P* (***θ***), we can apply linear stability analysis. That is we take the Jacobian of the left-hand side of Equation (1) to obtain,

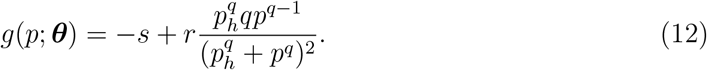

For a valid steady state *p*_∞_ ∈ *P* (***θ***), we can classify it as stable if sign(*g*(*p*_∞_; ***θ***)) = −1 and unstable if sign(*g*(*p*_∞_; ***θ***)) = 1, where we define

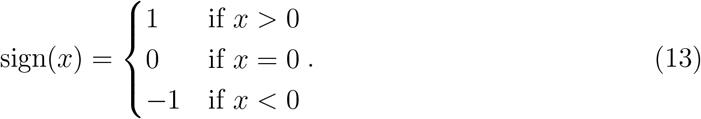

Given the steady state set, *P* (***θ***), and the Jacobian, *g*(*p*; ***θ***), are both functions of the model parameters ***θ***, we can apply profile-wise analysis. For an interest parameter, *ψ* (e.g., *ψ* = *c* to target the phosphorus loading as the interest parameter), let 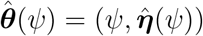 be a parameter set along the profile of *ψ* (Equation (10)). For any *ψ* within the profile-based confidence set (Equation (9)), then 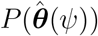 is the steady state set that within the confidence set and we can use 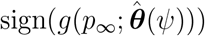 to determine the stability of within this profile-base confidence set. If both stable and bistable regimes exist within any confidence sets across all *ψ* ∈ *c, s, r, q, p*_*h*_, then the stability is considered non-identifiable.

Following form the profile-wise analysis of stability we can quantify profile-based confidence sets for tipping points. That is an equilibrium, *p*_∞_(*ψ*), is in the tipping point confidence if 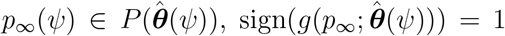, and *ψ* is in the confidence set of the profile likelihood targeting *ψ*. Thus we can sample *ψ* uniformly at random within the profile likelihood confidence set, and return every unstable equilibrium obtained via the profile-wise analysis. This provides a data-informed sensitivity analysis of the tipping point for each model parameter, and we can additional estimate an approximate confidence set for the tipping point using unions of the profile-based confidence sets.

### 2.6. Simulated data scenarios

Using the Carpenter model (Equation (1)) and the observation noise process (Equation (3)), we generate simulated dataset, representing a variety of environmental scenarios, to investigate whether the stability regime is practically identifiable given noisy observations similar to real lake monitoring data. In this manuscripte, we focus our attention on three simulated datasets that are produced under scenarios of the Carpenter model (Equation (1)) inspired by the study of Guttal and Jayaprakash (2007). These datasets are subsequently analysed using maximum likelihood estimation (MLE) and profile likelihood methods to estimate model parameters (Section 2.3), assess parameter identifiability (Section 2.4) and stability identifiability (Section 2.5), and preform sensitivity analysis of each parameter in relation to the tipping point (Section 2.5).

Each of three simulated data scenarios, represent the monitoring of a lakes recovery following an external disturbance event causing an influx of nurient load and increase in phosphorus concentration. Each scenario monitors the recovery over ten years using biannual measurements subject to a small amount of observation noise with a standard deviation of *σ* = 0.001 (g L^−1^). Simulated atasets are generated by numerically solving the Carpenter model (Equation (1)) using MATLAB’s `ode45` solver. Given an initial phosphorus concentration *p*(0) = *p*_0_, the model is numerically solved to obtain phosphorus concentrations at discrete time points *t*_0_ = 0, *t*_1_ = 0.5, …, *t*_20_ = 10 (years). The observed data are obtained by adding normally distributed measurement noise with zero mean and constant variance *σ*^2^ (Equation (3)) to the model output, accounting for measurement error and variability inherent in lake monitoring data.

As shown in Figure 6, each scenario represents a recovery towards an oligotrophic state. Scenario 1 (Figure 6(a)) and Scenario 3 (Figure 6(c)) produce time series that appear visually similar, displaying patterns almost identical and trends over time, despite their parameterisation and stability regimes being fundamentally different (Table 2). Scenario 1 (Figure 6(a)) is stable with a single oligotrophic equilibrium, whereas Scenario 3 (Figure 6(c)) is bistable and can transition between alternative stable states. Scenario 2 (Figure 6(b)) is also bistable, with a different parameterisation to Scenario 3 (Figure 6(c)), resulting in a tipping point that is very close to the peak phosphorus concentration (Table 2). Together, these three scenarios cover both stable and bistable regimes under oligotrophic conditions and provide an experimental framework for analyzing parameter estimation and identifiability. Although we have the ground truth parameters and stability regimes for the simulated data, our statistical framework does exploit this information as this is never available in reality. By using simulated data, however, it enables direct comparison between estimated and true system parameters and stability regimes.

**Table 2:**
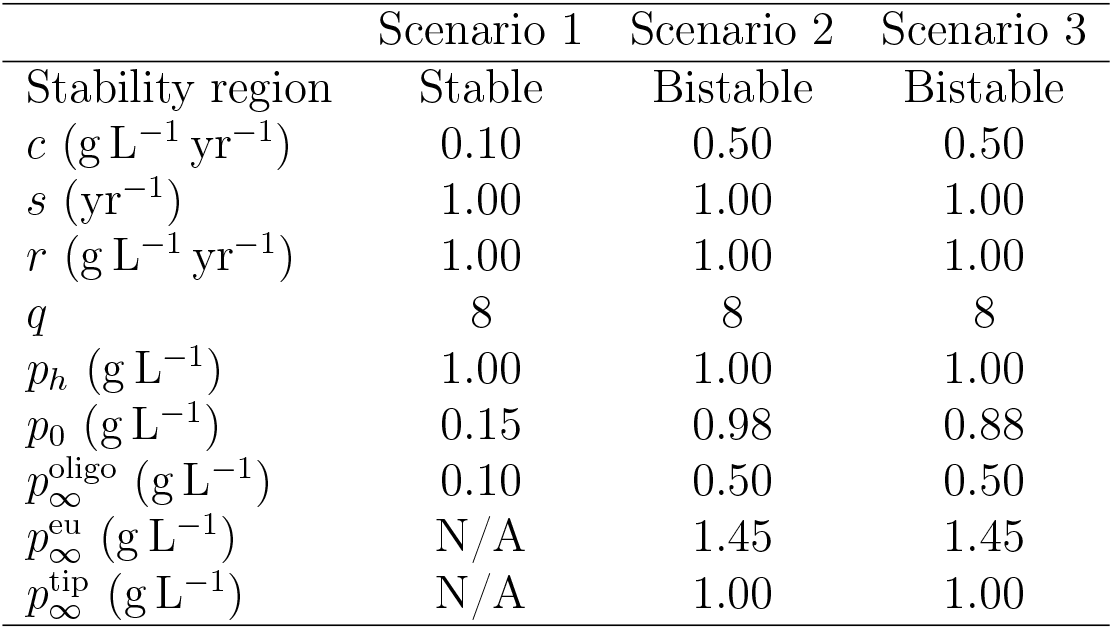
Parameter values and resulting equilibrium states for the three synthetic scenarios used in this study. Scenario 1 represents a stable oligotrophic system, while Scenarios 2 and 3 are bistable systems with both oligotrophic and eutrophic stable states. The table lists the external phosphorus loading rate (*c*), sink rate (*s*), maximum recycling rate (*r*), Hill coefficient (*q*), half-saturation constant (*p*_*h*_), initial condition (*p*_0_), and the long-term phosphorus concentrations at the oligotrophic 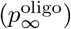, eutrophic 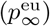, and unstable tipping point 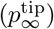 equilibria.

|  | Scenario 1 | Scenario 2 | Scenario 3 |
| --- | --- | --- | --- |
| Stability region | Stable | Bistable | Bistable |
| $c$ (g L<sup>-1</sup> yr<sup>-1</sup>) | 0.10 | 0.50 | 0.50 |
| $s$ (yr<sup>-1</sup>) | 1.00 | 1.00 | 1.00 |
| $r$ (g L<sup>-1</sup> yr<sup>-1</sup>) | 1.00 | 1.00 | 1.00 |
| $q$ | 8 | 8 | 8 |
| $p_h$ (g L<sup>-1</sup>) | 1.00 | 1.00 | 1.00 |
| $p_0$ (g L<sup>-1</sup>) | 0.15 | 0.98 | 0.88 |
| $p_\infty^{\text{oligo}}$ (g L<sup>-1</sup>) | 0.10 | 0.50 | 0.50 |
| $p_\infty^{\text{eu}}$ (g L<sup>-1</sup>) | N/A | 1.45 | 1.45 |
| $p_\infty^{\text{tip}}$ (g L<sup>-1</sup>) | N/A | 1.00 | 1.00 |

**Figure 6.**
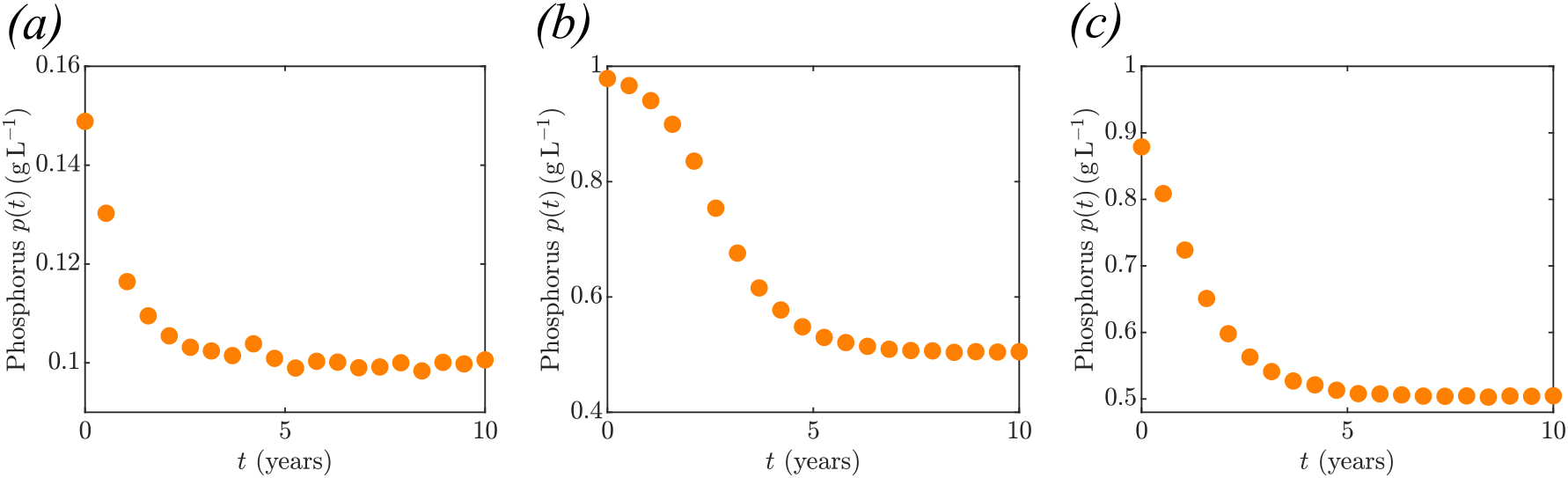
Simulated data scenarios representing lake monitoring data. Simulated phosphorus concentration measurements are taken using the Carpenter model (Equation (1)) with Gaussian observation noise (*σ* = 0.001 (g L^−1^)). Biannual observations are recorded at times *t*_0_ = 0, *t*_1_ = 0.5, …, *t*_20_ = 10 (years). Parameters for all three scenarios are given in Table 2. Scenario 1 (a) represents a stable system with a single stable oligotrophic equilibrium 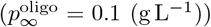. Scenario 2 (b) represents a bistable system with two stable equilibria, one oligotrophic 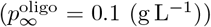 and one eutrophic 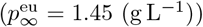, and one unstable tipping point 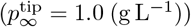. Scenario 3 (c) represent the same a bistable system as Scenario 2 (b) under different initial phosphorus concentrations. In (b) the initial phosphorus concentrations almost crossing the tipping point, whereas in (c) the initial phosphorus concentrations is well below the tipping point.

We focus on the oligotrophic recovery setting with low noise levels within this manuscript. However, the methods we present here are not restricted to this setting. We have considered eutrophication scenarios and others noise levels in Appendix C.2.

## 3. Results

To explore whether bistability can be identified from observational lake monitoring data, we analyse the three simulated data scenarios as described in Section 2.6. These datasets are analysed using the MLE and profile likelihood approaches to estimate model parameters (Section 2.3) and assess identifiability of parameters and stability regimes (Sections 2.4 and 2.5).

### 3.1. Parameter estimation using the MLE

For each simulated data scenario, the model parameters to be estimated, using the MLE (Section 2.3), are ***θ*** = (*c, s, r, q, p*_*h*_, *σ*), representing the phosphorus loading rate, phosphorus sink rate, phosphorus recycling rate, Hill coefficient, the phosphorus concentration at half-maximum recycling, and the noise level, respectively (Table 1).

The MLEs and the corresponding 95% Wald confidence intervals for the model parameters in the three main scenarios are presented in Table 3.

**Table 3:**
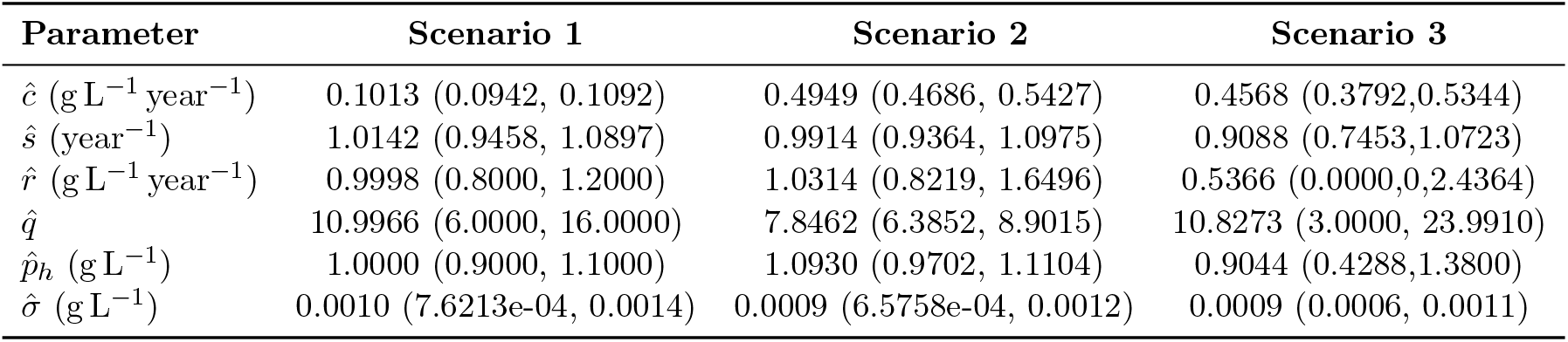
Maximum likelihood estimates and 95% Wald style confidence intervals for the three oligotrophic lake stability regimes: Section 2.6 and Figure 6 for a description of simulation Scenarios.

| Parameter | Scenario 1 | Scenario 2 | Scenario 3 |
| --- | --- | --- | --- |
| $\hat{c}$ (g L<sup>-1</sup> year<sup>-1</sup>) | 0.1013 (0.0942, 0.1092) | 0.4949 (0.4686, 0.5427) | 0.4568 (0.3792, 0.5344) |
| $\hat{s}$ (year<sup>-1</sup>) | 1.0142 (0.9458, 1.0897) | 0.9914 (0.9364, 1.0975) | 0.9088 (0.7453, 1.0723) |
| $\hat{r}$ (g L<sup>-1</sup> year<sup>-1</sup>) | 0.9998 (0.8000, 1.2000) | 1.0314 (0.8219, 1.6496) | 0.5366 (0.0000, 2.4364) |
| $\hat{q}$ | 10.9966 (6.0000, 16.0000) | 7.8462 (6.3852, 8.9015) | 10.8273 (3.0000, 23.9910) |
| $\hat{p}_h$ (g L<sup>-1</sup>) | 1.0000 (0.9000, 1.1000) | 1.0930 (0.9702, 1.1104) | 0.9044 (0.4288, 1.3800) |
| $\hat{\sigma}$ (g L<sup>-1</sup>) | 0.0010 (7.6213e-04, 0.0014) | 0.0009 (6.5758e-04, 0.0012) | 0.0009 (0.0006, 0.0011) |

Figure 7 (a)–(c) shows that the Carpenter model (Equation (1)) parameterised by the MLE (Table 3) provides a close fit to the simulated phosphorus data across oligotrophic regimes, both stable and bistable. This is demonstrated by comparing the model trajectories (blue) using the MLE parameters (Table 3) with the simulated phosphorus concentrations (orange circles) generated for each Scenario (Figure 6).

**Figure 7.**
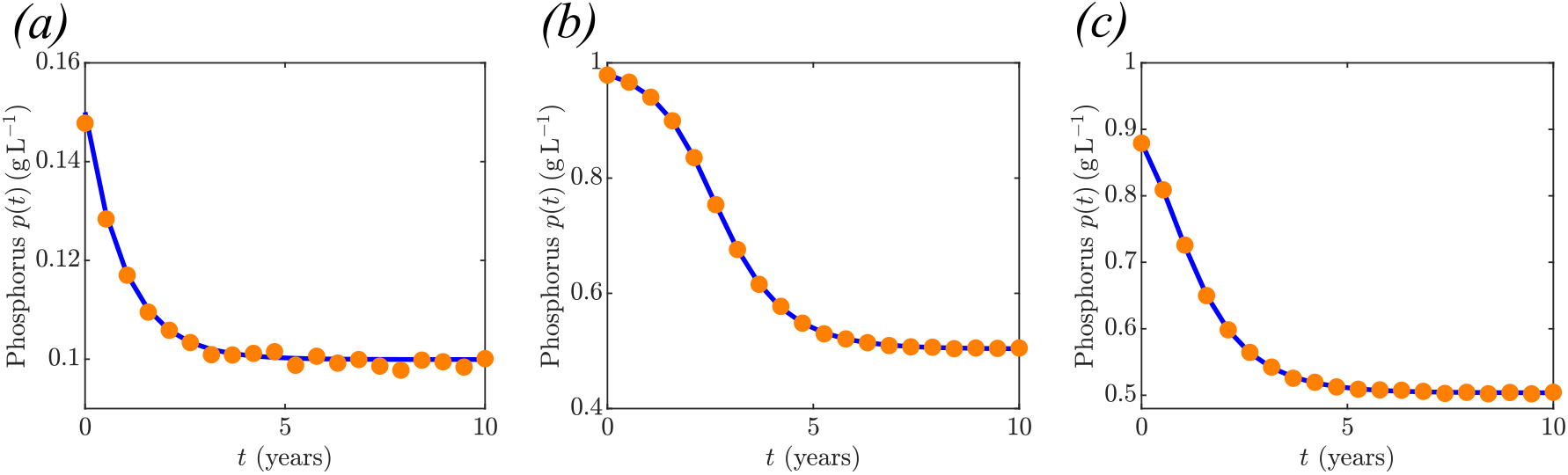
Model fit at the MLE for the three simulated data scenraios provided in Section 2.6 (Figure 6). Simulated phosphorus concentration observations (orange circles) are shown alongside the MLE trajectory (blue line), representing the expected mean phosphorus concentration under the parameter estimates provided in Table 3.

Through the invaraince property of the MLE, we can numerically obtain the MLE for the equilibria in each scenario given the MLE for the parameters (Table 3). In Scenario 1, the estimated parameters produce a single stable equilibrium corresponding to a long-term oligotrophic phosphorus concentration of 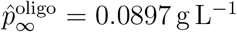, which corresponds well with the ground truth solution, 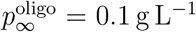 (Table 2). This confirms that, under low noise conditions, the estimation procedure accurately recovers the system’s dynamics. Similarly, for Scenario 2 the MLE leads to two stable equilibria: an oligotrophic equilibrium at 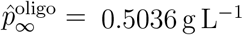 and a eutrophic equilibrium at 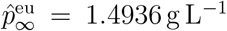. These estimated stable states are closely aligned with the ground truth equilibria of Scenario 2, 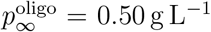 and 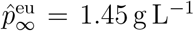 (Table 2). In contrast, the MLE obtained for Scenario 3 incorrectly leads to a stable system, 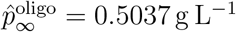.

### 3.2. Parameter identifiability analysis

Using the structural identifiability analysis software GenSSI (Chiş et al., 2011), we conclude that the Carpenter model is only locally structurally identifiable. This means that each parameter can be uniquely determined within a neighbourhood of the true parameter values given the model structure. The implication is that while we can estimate unique parameter values within a small range around the true values, the model does not guarantee a unique solution across the entire parameter range. As a result, some parameters may be practically identifiable with real data, while others may remain non-identifiable due to limitations in data and noise. To explore this further, we conduct a practical identifiability analysis based on profile likelihoods (Section 2.4). Results are presented in Figures 8–10.

**Figure 8.**
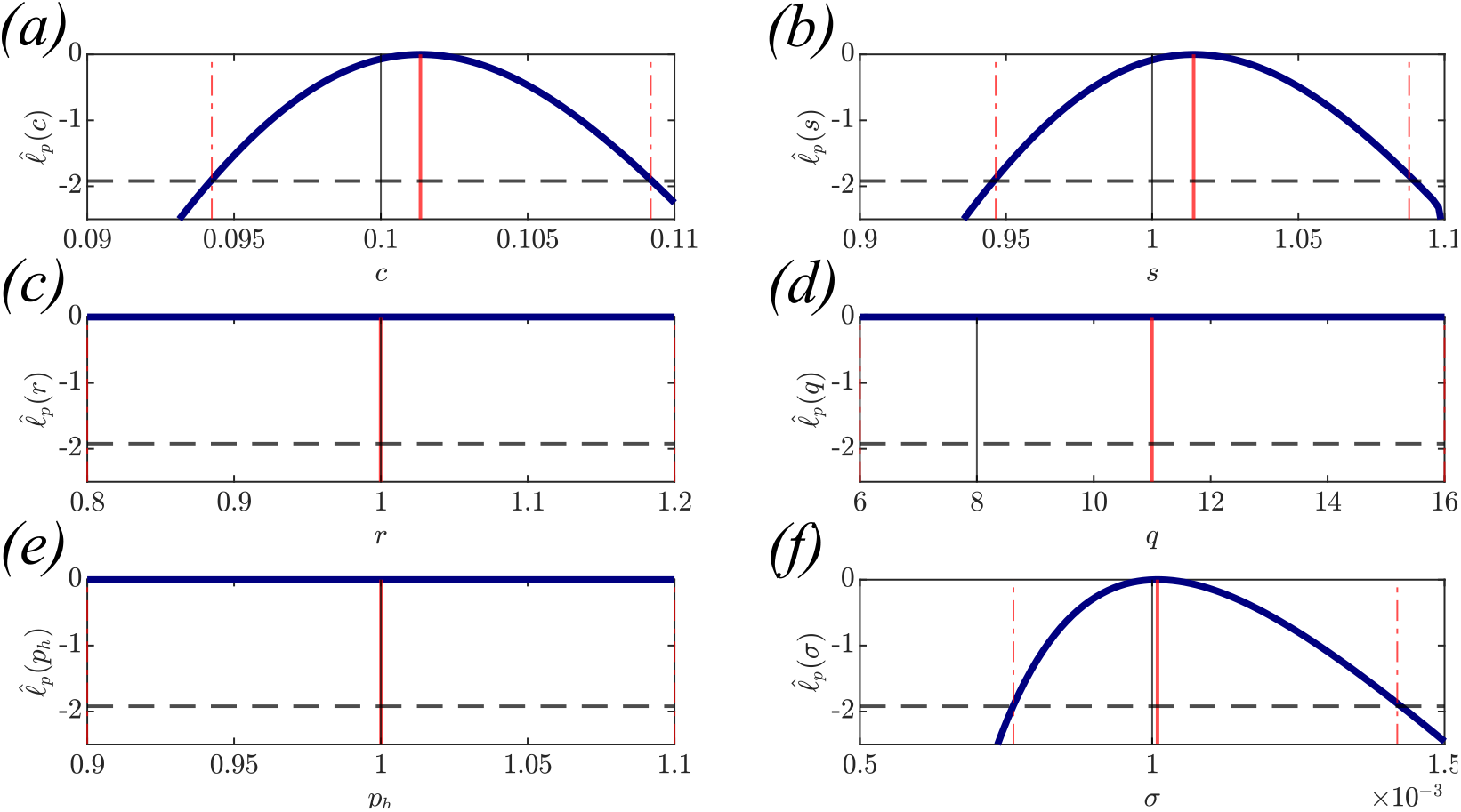
Profile likelihood plots for the Scenario 1 (Figure 6(a)). Each plot shows the univariate profile likelihood curve (solid blue line), the MLE (red solid line) for each parameter, the ground truth parameter values (black solid line), and the 95% confidence intervals (red dashed lines) based on Wilks’ theorem with threshold of 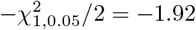 (black dashed line). Ground truth parameters can be found in Table 2 and MLE values in Table 3.

The analysis of Scenario 1 is shown in Figure 8, we see that model parameters *c, s*, and *σ* (Figure 8(a)–(b),(f)) are identifiable as indicated by well-defined peaks and narrow profile-based confidence intervals. However, the parameters that contribute the nonlinear phosphorus recycling term, *r, q*, and *p*_*h*_ (Figure 8(c)–(e)) are practically non-identifiable with the flat profile curves. This is intuitively reasonable, since Scenario 1 represents a stable oligotrophic system, and this can only occur if the loading and sink terms dominate. As a result, there is little information on this term contained in the simulated data.

Figure 9 presents the profile likelihood results for Scenario 2. Here, all model parameters are identifiable, as within well-defined and narrow profile likelihood curves shown in Figure 9. This contrasts with the poor identifiability observed in Scenario 1 cases (Figure 8), where the profile likelihood curves in the figure show broad, flat regions indicating that a wide range of parameter values fit the data equally well. However, Figure 9 shows sharply peaked and narrow profile likelihood curves, demonstrating that the likelihood is very sensitive to changes in any of parameter values, including the nonlinear recycling rate parameters *r, q*, and *p*_*h*_ (Figure 9(c)–(e)). These well-defined peaks correspond to tighter confidence intervals.

**Figure 9.**
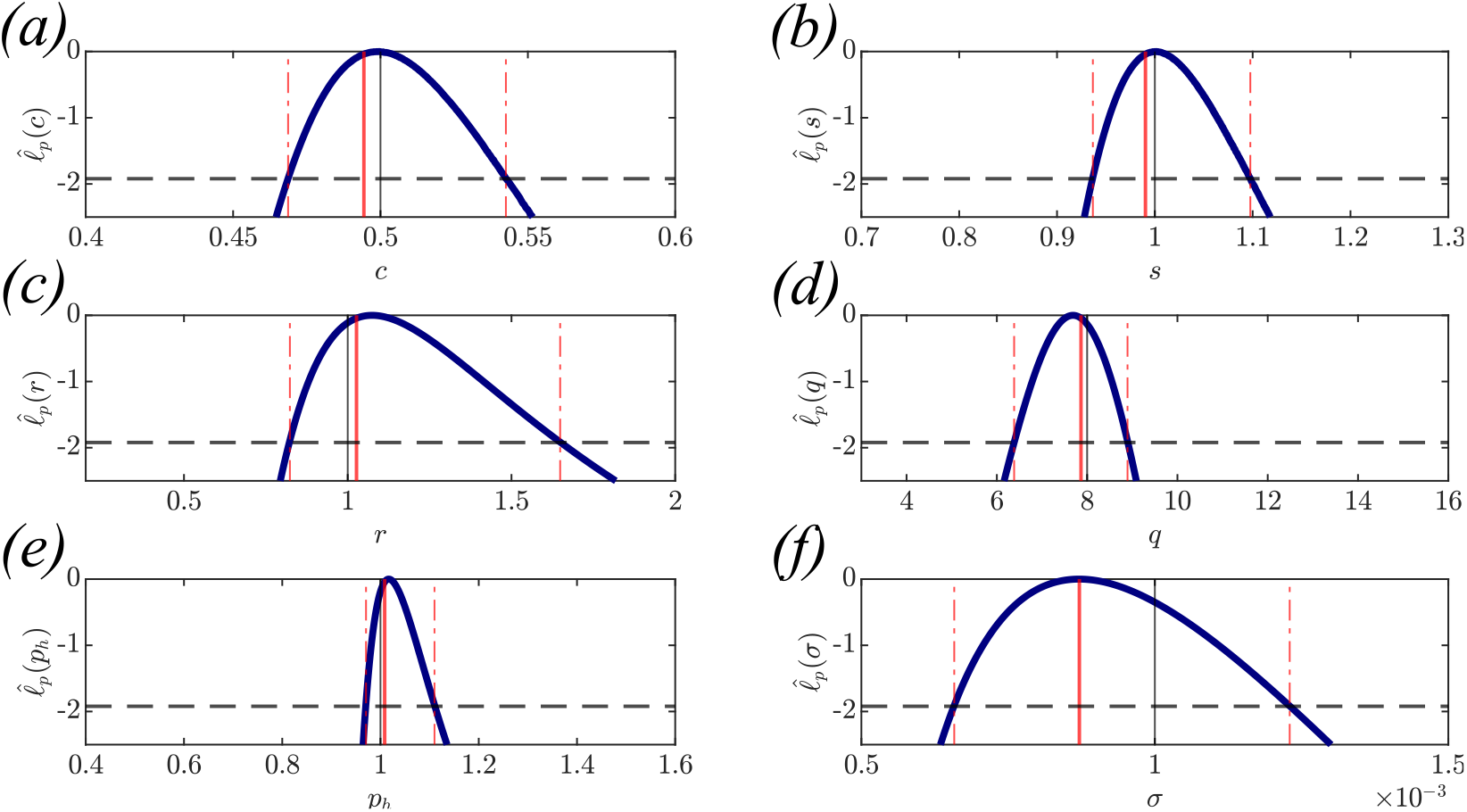
Profile likelihood plots for the Scenario 2 (Figure 6(b)). Each plot shows the univariate profile likelihood curve (solid blue line), the MLE (red solid line) for each parameter, the ground truth parameter values (black solid line), and the 95% confidence intervals (red dashed lines) based on Wilks’ theorem with threshold of 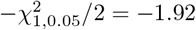 (black dashed line). Ground truth parameters can be found in Table 2 and MLE values in Table 3.

For Scenario 3 (Figure 10), the parameters *c, s, p*_*h*_, and *σ* (Figure 10(a)–(b),(e)–(f)) have narrow and well-defined peaks, indicating that these parameters are practically identifiable. However, two of the non-linear recycling term paremeters *r* and *q* (Figure 10(c)–(d)) are only partially identifiable. While a lower confidence interval in both cases is obtained, the likelihood profiles appear to be monotonically increasing.

**Figure 10.**
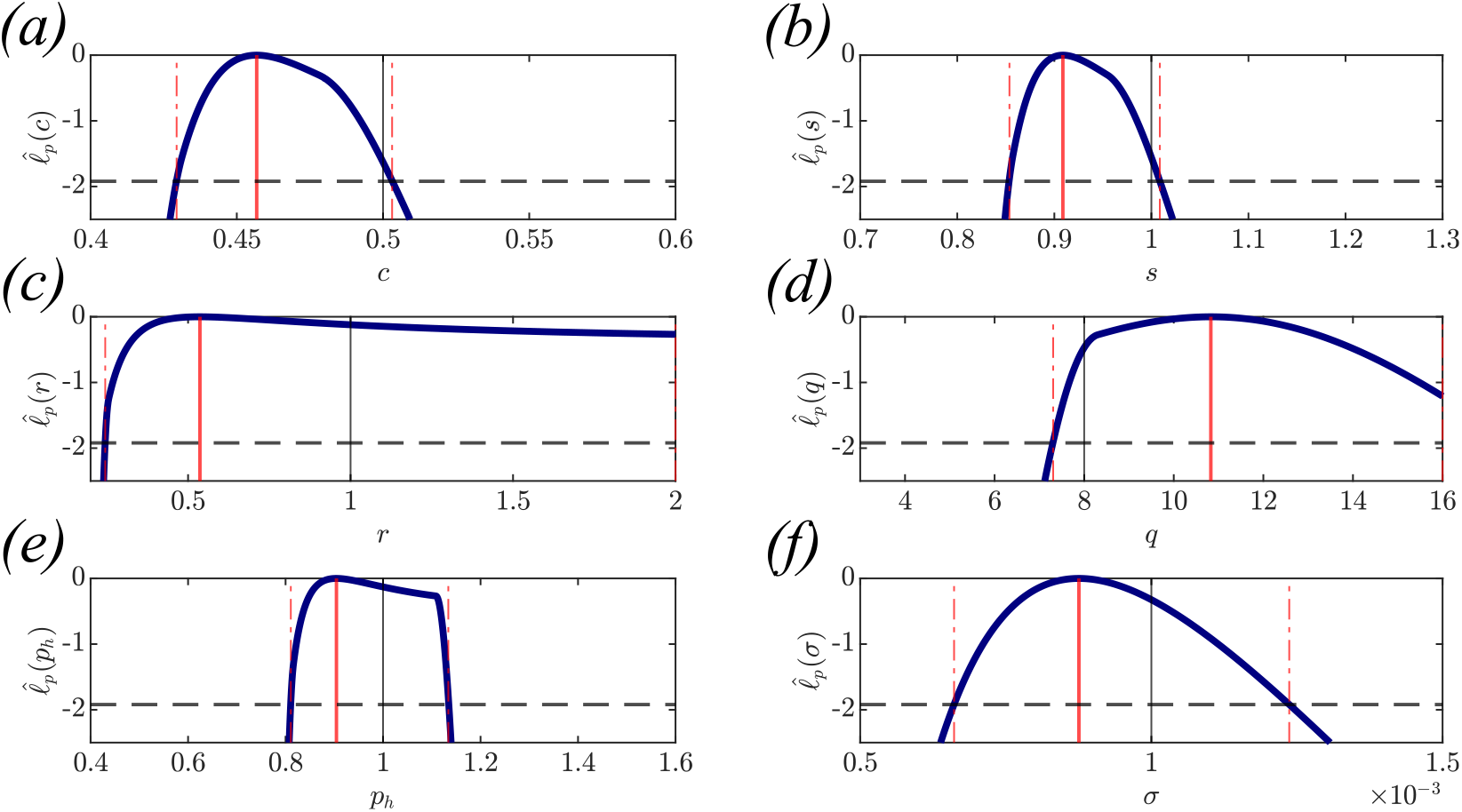
Profile likelihood plots for the Scenario 3 (Figure 6(c)). Each plot shows the univariate profile likelihood curve (solid blue line), the MLE (red solid line) for each parameter, the ground truth parameter values (black solid line), and the 95% confidence intervals (red dashed lines) based on Wilks’ theorem with threshold of 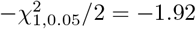 (black dashed line). Ground truth parameters can be found in Table 2 and MLE values in Table 3.

### 3.3. Stability identifiability and tipping-point sensitivity

The results in Section 3.2 show that, depending on the ground truth system, parameter identifiability is not guaranteed. However, the profiles alone do not provide direct insight into the identifiability of system stability or sensitivity of tipping points, if they are present. For this we use our application of profile-wise analysis presented in Section 2.5.

For the three simulated data scenarios (Figure 6), we apply the profile-wise analysis of stability. Within the profile-based 95% CI for each interest parameter, the profile is tracked along the grid (Equation 10). For each point, the stability is evaluated (using Equations (11)– (13)) and visualised on the profile by indicating stable systems as blue and bistable as red. Figure 11, presents the results across all three scenarios.

**Figure 11.**
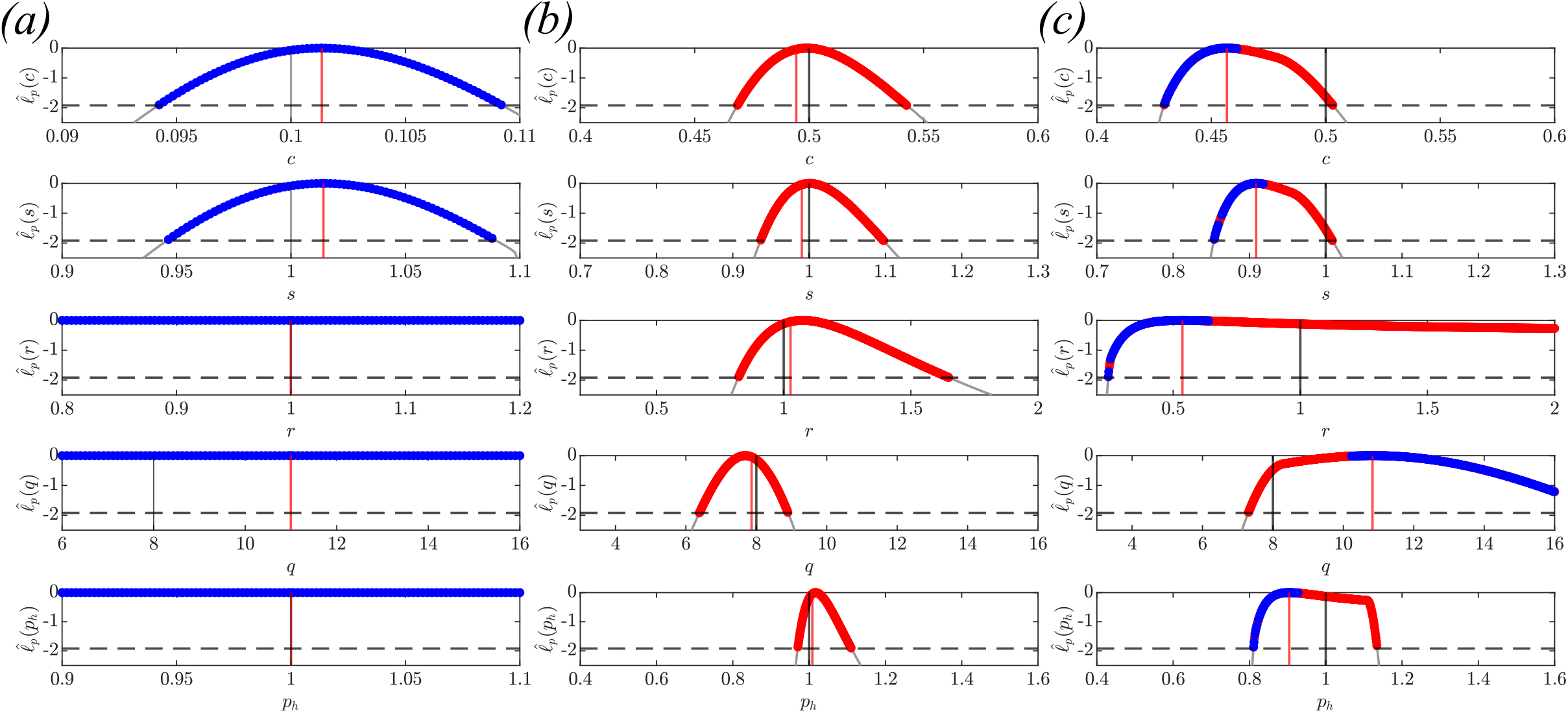
Visualisation of profile-wise identifiability analysis for stability classification across the three simulated data scenarios: (a) Scenario 1, (b) Scenario 2, and (c) Scenario 3. Each plot profile likelihood curves, as given in Figures 8–10, with grid points colour coded based on stability analysis (stable = blue, bistable = red). Also indicated are the MLE (red solid line) for each parameter, the ground truth parameter values (black solid line), and the threshold 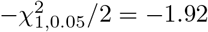 (black dashed line) for the 95% confidence interval under Wilks’ Theorem. Ground truth parameters can be found in Table 2 and MLE values in Table 3.

For both Scenario 1 and 2, the stability is identifiable, with the profile-wise stability analysis leading to exclusively stable systems (Scenario 1; Figure 11(a)), or exclusively bistable systems (Scenario 2; Figure 11(b)). In contrast, the profile-wise stability analysis of Scenario 3 exhibits cases of stable systems and bistable systems with the profile-based confidence intervals (Figure 11(c)), we conclude that stability is non-identifiable for Scenario 3. These results indicate that parameter non-identifiability does not imply stability non-identifiability (Figure 11(a)). However, for the bistable scenarios, it seems that identifiability of stability is related to parameter identifiability. We discuss the implications of this for management in Section 4.

Following from the results in Figure 11, we evaluate the sensitivity in the tipping point estimates to changes in the parameters along the profile. That this, we examine how tipping points are affected by data-driven changes in model parameters (Liu et al., 2026). We can further take the union of profile-wise 95% confidence intervals for the tipping point across all interest parameter variations to obtain an approximation to the over all 95% confidence intervals for the tipping point.

Table 4 present the profile-wise sensitivity analysis for both Scenarios 2 and 3 correspond to parameter sets within the bistable regime. Overall, we note that the 95% confidence intervals for the tipping point estimate are much narrower for Scenario 2 compared with Scenario 3. In terms of sensitivity, we see that profile-wise changes in recycling rate, *r*, and the half-saturation parameter, *p*_*h*_, have the greatest impact on the uncertainty in the tipping point estimate. These findings underscore the pivotal role of specific parameters, particularly *p*_*h*_ and *r*, in determining the positions of tipping points while man have management implications (Section 4).

**Table 4:** Profile-wise sensitivity of the tipping point estimate, 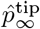, to changes in the various models parameters along the univariate profile curve (as per Section 2.5) for both Scenarios exhibiting bistablity (Figure 11(b)–(c)). Overall estimate uncertainty is calculated using the union of all profile-wise confidence intervals and the width of this interval is reported.

| Parameter | Scenario 2<br>$\hat{p}_\infty^{\text{tip}}$ (95% CI) | Scenario 3<br>$\hat{p}_\infty^{\text{tip}}$ (95% CI) |
| --- | --- | --- |
| $c$ | [0.9957, 1.0033] g L<sup>-1</sup> | [0.8876, 1.0737] g L<sup>-1</sup> |
| $s$ | [0.9955, 1.0034] g L<sup>-1</sup> | [0.8874, 1.0732] g L<sup>-1</sup> |
| $r$ | [0.9948, 1.0049] g L<sup>-1</sup> | [0.8664, 1.0738] g L<sup>-1</sup> |
| $q$ | [0.9951, 1.0046] g L<sup>-1</sup> | [0.8802, 1.0730] g L<sup>-1</sup> |
| $p_h$ | [0.9948, 1.0052] g L<sup>-1</sup> | [0.8659, 1.0714] g L<sup>-1</sup> |
| Union | [0.9948, 1.0052] g L<sup>-1</sup> | [0.8659, 1.0738] g L<sup>-1</sup> |
| CI width | 0.0104 g L<sup>-1</sup> | 0.2079 g L<sup>-1</sup> |

## 4. Discussion

In this work, we assess how informative standard ecological monitoring data are for reliable parameter estimation and determination of the ecosystem exhibits bistable behaviour. Using a classical nonlinear dynamical model of lake eutrophication, we explore how practical parameter identifiability impacts the identifiability of system stability. Our scenario analysis shows that identifying stability regime using standard monitoring data is challenging, as identifiable parameters may not be sufficient to ensure the stability regime can be determined. Conversely, non-identifiable parameters do not necessarily lead to non-identifiability of the stability regime. A key finding is, for a bistable system, that parameter and stability identifiability improve when data is collected near the tipping point. These findings have broad implications for modelling regime shifts in ecosystems such as lakes, coral reefs, or forests, where transitions between alternative stable states are determined by the parameter uncertainty. Even when parameter estimates are well constrained, the qualitative system behaviour may remain unclear, highlighting the limits of typical monitoring data for determining ecosystem stability.

We use a well-known lake eutrophication model (Equation (1)) as a canonical example of bistable ecosystems. We focused on oligotrophic lake regimes under three main scenarios to identify whether the system, is capable of transition between oligotrophic and eutrophic states or remains solely oligotrophic regardless of the external disturbances. We use observed data and tools from parameter identifiability analysis, such as likelihood profiles (Section2.4) and profile-wise prediction (Section2.5). Parameter identifiability is widely studied in both deterministic and stochastic biological models, using a range of techniques including frequentist methods such as profile likelihood analysis (Simpson et al., 2022; Warne et al., 2024; Raue et al., 2009), as well as Bayesian approaches (Browning et al., 2020; Hines et al., 2014). Similarly, several studies have examined the bistability of the Carpenter model under noisy data conditions (Guttal and Jayaprakash, 2007; Carpenter et al., 1999) leading to the risk of irreversible degradation indicator (Wu et al., 2019). However, these methods for detecting ecological risk do not consider the identifiability of the ecosystem stability. A key question in this study is whether the lake monitoring data contain enough information to distinguish between stable and bistable regimes. To the best of our knowledge, no existing studies have specifically applied parameter identifiability (Section 2.4) to the bistable ecosystems using profile likelihood analysis. Our work addresses this gap by examining whether observed data contain sufficient information to estimate parameters and determine stability across different scenarios.

In terms of parameter identifiability, presence of flat regions in the profile likelihood analysis of Scenario 1, a stable system, indicates practical non-identifiability for the parameters related to recycling *r, q*, and *p*_*h*_ (Figure 8). In such cases, even though the MLE is obtained (see Table 3), any parameter combination within the confidence region can represent the true system. For Scenarios 2 and 3 (Figures 9 and 10), both bistable systems, the recycling parameters can be identifiable or partially identifiable. A key reason for this reduced identifiability in Scenario 3 is the choice of initial condition, representing the peak phosphorus following a disturbance. Since the system never gets close enough to the tipping point, it does not exhibit the more complex nonlinear effect of the recycling term that starts to dominate close to the tipping point. This reduces the information available in the data about *r* and *q*. Therefore, starting the system closer to the tipping pointm increases its sensitivity and improves the estimation of the recycling parameters more accurately.

This parameter identifiability analysis alone is not sufficient to determine stability identifiability. For example, in Scenarion 1 (Figure 8), independently varying the values of the non-identifiable parameters, while keeping the identifiable parameters fixed, could results in a stable system or a bistable system within the same univariate confidence regions (See Appendix C.1). From a management perspective, this illustrates a substantial risk in that a system may appear to be safely operating within a stable regime, it may be bistable an capable of crossing a tipping point toward a eutrophic state. This demonstrates how practical non-identifiability can hide the true stability structure of the ecosystem. The lake may appear stable based on the estimated MLE alone, but the real system behaviour could be much more sensitive than it seems. This misclassification directly affects management because decisions made under the assumption of a stable regime increase the chance of a critical transition, increasing the risk of unexpected regime shifts in lake ecosystems.

Our findings highlight the key points in applying identifiability analysis to detect the stability of the bistable ecosystem. Even when a model has identifiable parameters (Figure 11b), it does not guarantee that the system’s qualitative behavior, such as stability (Figure 11c), is also identifiable from data. In contrast, non-identifiability of certain parameters (Figure 8) does not imply that stability is non-identifiable (Figure 11a). The stability of the system seems to become identifiable only when the system is close to the tipping point (Figure 6b). These results demonstrate that even well-constrained parameter estimates may not fully reveal the system’s stability, emphasizing the importance of evaluating stability directly rather than assuming it follows from parameter identifiability. Moreover, when we apply this framework at higher noise levels (*σ* = 0.1 g L^−1^), it increases the uncertainty in identifying the stability classification of the ground truth for both oligotrophic and eutrophic conditions (Appendix C.3).

While this work provides key understanding relevant to the management of ecosystems, several limitations should be addressed in future work. First, we only consider the Carpenter model (Equation (1)), which is ideal for studying the bistability of a lake rather than other models (Chang et al., 2022), and it is a commonly used example for lake eutrophication (Wu et al., 2019; Guttal and Jayaprakash, 2007; Carpenter et al., 1999). However, the statistical analysis and identifiability methods we use are completely general and can be applied to other nonlinear ecological or biological systems. Future studies should test whether similar identifiability patterns occur in other bistable systems to assess the generality of our conclusions.

Secondly, our results show that, under bistable conditions, the deterministic model does not consistently recover the correct stability regime; in some cases it correctly identifies bistability, while in others it fails to do so. This highlights the need for alternative way to improve the reliability of regime identification. This approach aligns with the findings of Guttal and Jayaprakash (2007) that intrinsic noise can enhance the detectability of critical transitions. Therefore, extending the current analysis to the stochastic version (Wang and Qi, 2018), as suggested by Browning et al. (2020), could provide a deeper understanding of the regime of the bistable ecosystem. The stochastic models the likelihood may be intractable; however, there are statistical methodologies available to handle this setting (Warne et al., 2024).

We identify that changes in some parameters within their uncertainty ranges can lead to changes in the system’s stability (Appendix C.1). This means some parameters have a strong influence on stability identification, while others have a minor effect. Therefore, it is important to determine which parameters require more attention and more accurate estimation. Building on this finding, future work should incorporate optimal experimental design methods to identify the most informative sampling strategies that maximize parameter and stability identifiability (Warne et al., 2017).

In conclusion, our findings highlight the limitations of classical statistical methods, such as maximum likelihood estimation and standard model fitting, in accurately quantifying stability uncertainty. While estimated system trajectories may indicate general system states, key parameters remain non-identifiable (Figure 8), highlighting the need for dedicated identifiability analysis. Our results suggest that bistability is only partially identifiable using standard monitoring data. In some cases, system stability can be estimated when the initial condition is close to the tipping points (Figure 11 (b)). Addressing this challenge requires improved monitoring designs that provide data informative enough to determine both parameter estimates and system stability.

From a management perspective, the framework presents a step beyond fitting system dynamics by examining uncertainty in stability regimes. We make a novel contribution by extending profile likelihood analysis to study ecosystem stability and quantify the uncertainty surrounding bistability. Previous approaches, such as risk-based indicators (Wu et al., 2019), can signal ecological degradation but haven’t determined whether alternative stable states exist or whether a tipping point can be identified without the system crossing it. By quantifying the strength of evidence for alternative stable states, our approach offers a practical tool for risk-aware management of bistable ecosystems. This approach is important because misidentifying stability regimes can lead to harmful management decisions in ecosystems, where small perturbations may trigger irreversible shifts. Our method provide, accurate identifiability analysis that could be critical for responsible conservation planning.

## Authorship contribution statement

**Dasuni Amanda Salpadoru:** Writing – original draft, Visualization, Validation, Software, Methodology, Formal Analysis. **Matthew P. Adams:** Writing – review & editing, Validation, Supervision, Conceptualization. **Kate Helmstedt:** Writing – review & editing, Supervision, Conceptualization. **David J. Warne:** Writing – review & editing, Validation, Software, Methodology, Formal Analysis, Supervision, Conceptualization.

## Software and data availability section

The MATLAB software and associated data will be made available on GitHub.

## Acknowledgments

DAS is supported by a Research Training Program Scholarship. DJW is supported by an Australian Research Council (ARC) Early Career Researcher Award (DE250100396). DJW acknowledges continued support from the Centre for Data Science at the Queensland University of Technology. MPA and KJH acknowledge funding from the ARC SRIEAS Grant Securing Antarctica’s Environmental Future (SAEF; SR200100005).

## Appendix A. Stability analysis

As discussed in the stability analysis (Section 2.2), the bifurcation diagram changes (Figure 3) significantly as the nutrient loading rate *c* varies. In addition to variations in *c*, changes in the Hill coefficient *q* also play a critical role in shaping the stability structure of the Carpenter model (Equation (1)). To study the effects of *q*, we fix all other parameters in the model to biologically reasonable values based on Guttal and Jayaprakash (2007). We set the phosphorus sink rate *s* = 1 year^−1^, the maximum recycling rate *r* = 1 gL^−1^year^−1^, and the phosphorus concentration at which recycling reaches half of its maximum, *p*_*h*_ = 1 gL^−1^.

To understand how changes in the Hill coefficient *q* influence the bifurcation diagram, we analyze the steady-state behavior of the Carpenter model (Equation (1)) by substituting the fixed parameter values *s* = 1 year^−1^, *r* = 1 gL^−1^year^−1^, and *p*_*h*_ = 1 gL^−1^, which simplifies the model to

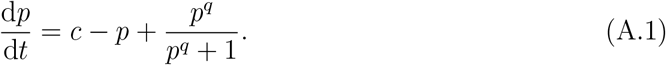

At steady state, we set 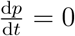, giving a simplified version of Equations (2) and (11)

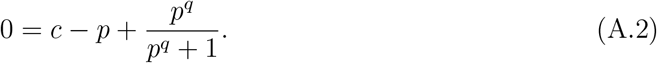

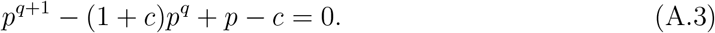

For positive integer values of *q*, Equation (A.3) is a polynomial of degree *q* + 1 in *p*, and the number of real roots it possesses depends on the value of *q*. For low values of *q*, the equation produces a single real root, while for higher *q*, the system may exhibit three real roots, showing bistability (Figure A.1).

**Figure A.1.**
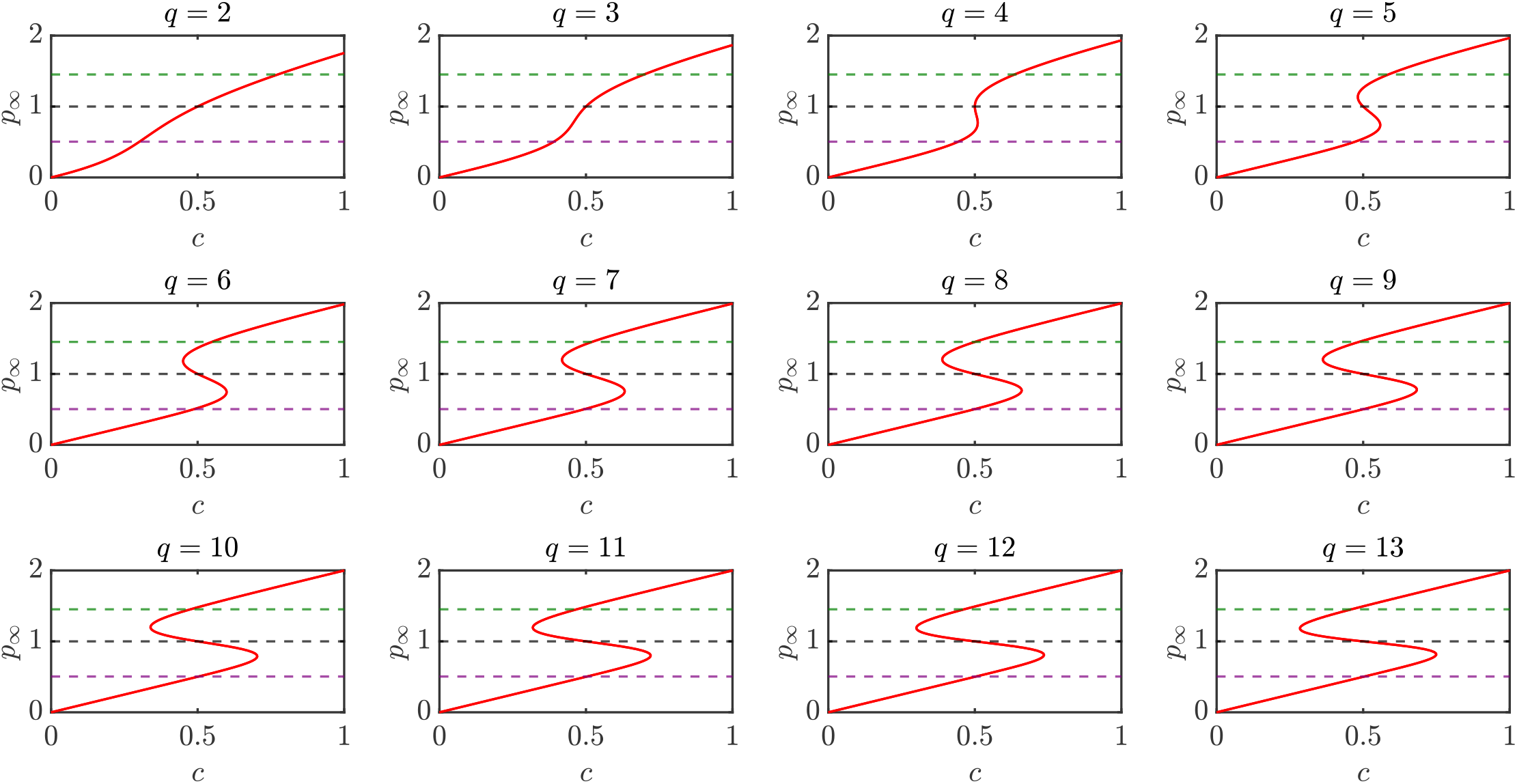
Effect of the Hill coefficient *q* on the steady-state phosphorus concentration *p*_∞_, given by the solutions to the steady-state polynomial equation. Each subplot corresponds to a fixed value of *q*, ranging from 2 to 13, with the nutrient loading rate *c* varying over the interval [0, 1] g L^−1^ year^−1^. The red curve in each plot represents the set of phosphorus concentrations *p* that satisfy the equilibrium condition 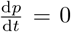 for a given *c*. Dashed horizontal lines indicate key reference points: the oligotrophic stable equilibrium at 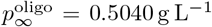, the eutrophic stable equilibrium at 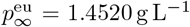, and the critical tipping point at 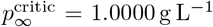. The study considers the reference value as *q* = 8. When the red contour is single-valued with respect to *c*, the system is non bistable. When the red contour folds and shows multiple values of *p* for the same *c*, the system becomes bistable, showing two stable equilibria and one unstable equilibrium.

## Appendix B. Confidence Interval Calculation

In this section, we discuss standard approaches for constructing confidence interval: the Wald method based on the observed Fisher Information Matrix (FIM) and profile likelihood method based on Wilk’s theorem.

### Appendix B.1. Wald-style Confidence Interval (Fisher Information-Based)

The Wald confidence interval is derived under the assumption that the maximum likelihood estimator (MLE), 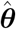, is approximately normally distributed in large samples. It is based on the result that 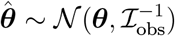, where ℐ_obs_ is the observed Fisher Information Matrix, defined as the negative of the second derivative (Hessian) of the log-likelihood function compute at the MLE. The confidence interval for each parameter ***θ***_*i*_ is given by

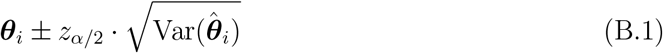

where the variance is extracted from the diagonal of 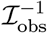, and *z*_*α/*2_ is the standard normal quantile (1.96 for a 95% CI).

The Wald method is computationally efficient and widely used because it is easy to use in models that are defined using parameters. However, its reliability depends on the asymptotic normality of the estimator. It performs poorly in small samples, near boundary values, or when the parameter is weakly or nonidentifiable. In such cases, the likelihood may not look like a smooth, bell-shaped curve, breaking the assumption that the maximum likelihood estimator is approximately normally distributed. Therefore, Wald intervals are best suited for identifiable parameters in large-sample settings (Casella and Berger, 2002; Agresti and Caffo, 2000).

#### Appendix B.2. Profile Likelihood Confidence Interval (Wilks’ Theorem-Based)

Confidence intervals can be estimated from likelihood profiles by using theory based on Wilks’ theorem are describing the asymptotic behaviour of the from the likelihood ratio test statistic used for model selection with nested models (Pawitan, 2001; Wilks, 1938). Wilks’ theorem states that, under regularity conditions, the likelihood ratio statistic

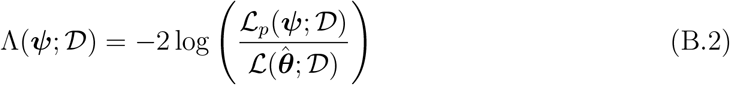

where 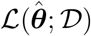 is the maximum likelihood under the full model for a given data set 𝒟, and ℒ_*p*_(***ψ***; 𝒟) is the profile likelihood for the parameter of interest ***ψ***, obtained by fixing ***ψ*** and maximizing the likelihood over the nuisance parameters. By simplifying, the expression becomes

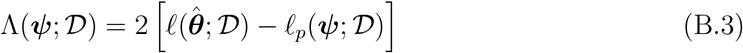

where 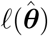 and *ℓ*_*p*_(***ψ***) denote the log-likelihood and profile log-likelihood, respectively. Wilks showed that, asymptotically, Λ(***ψ***; 𝒟) is distributed according to a chi-squared distribution with degrees of freedom equal to the number of tested parameters (that is the dimension of the interest parameter *d*_***ψ***_). Here, 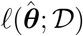 is the maximum log-likelihood, and 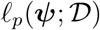 is the profile log-likelihood for the parameter of interest ***ψ***. The confidence interval for ***ψ*** consists of all values satisfying

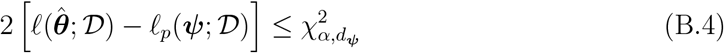

where 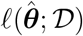 is the maximum log-likelihood value, and 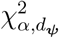 is the chi-square critical value at significance level *α* and degrees-of-freedom *d*_***ψ***_ corresponding to the dimensionality of the interest parameter. Once the threshold is calculated, a value of − 1.92 corresponds to an approximate 95% confidence interval for the univariate profile likelihoods. Finally, visual representations can be used to further assess the identifiability of each parameter. By examining the shape of the profile likelihood curves, we can determine whether the model parameters are identifiable or not (Figure 5).

In contrast to the Wald approach, this method does not depend on normality assumptions. It is therefore well-suited for nonlinear or poorly identifiable models, where the likelihood surface may be skewed or flat. The profile likelihood method does not depend on normality, making it suitable for models where the likelihood surface is asymmetric or parameters are weakly identifiable. However, they are computationally intensive, as each point in the profile requires an optimization over nuisance parameters. However, due to their flexibility, this is valuable for uncertainty quantification in ecological and biological modeling (Wilks, 1938; Raue et al., 2009).

#### Appendix B.3. Numerical Approximation to the Observed Fisher Information Matrix

To quantify uncertainty in the parameter estimates of the Carpenter model (Equation (1)), we compute confidence intervals using the Observed Fishery Information matrix (FIM) (Equation (B.6)). This approach depends on approximating the second derivative (Hessian matrix) of the log-likelihood function evaluated at the maximum likelihood estimate (MLE).

In the classical approach, the variance of the MLE is estimated using the Fisher Information Matrix (FIM), which quantifies the second derivative of the log-likelihood function around the MLE,

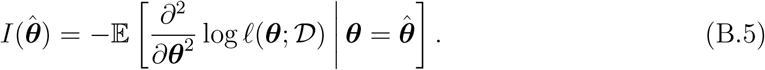

Here, 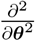 denotes the Hessian of the log-likelihood with respect to the parameters. The expectation with respect to the data, 𝒟, is taken as negative because the second derivative of the log-likelihood is negative at the maximum. In most practical applications, the exact expectation is difficult to obtain. Therefore, observed FIM is used to calculate the confidence interval of model parameters,

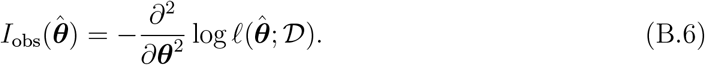

This directly calculates the second derivatives of the log-likelihood at the MLE 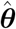, for the observed data, rather than the expectation of the data. However, building the Hessian matrix analytically is intractable for nonlinear models. Therefore, in this study, we use the second derivatives that are numerically approximated using a central finite difference method (Appendix A). It is used because it provides a more accurate approximation of second derivatives, with smaller error compared to forward or backward difference methods.

Once the observed FIM (Equation (B.6)) has been computed, it provides an estimate of the asymptotic covariance matrix of the MLE. Using this approximation, the Wald confidence ellipsoid can be approximated asymptotically using 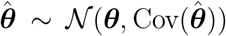 where 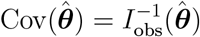.

The observed FIM (Equation (B.6)) is defined as the negative Hessian (the matrix of second-order partial derivatives) of the log-likelihood function. However, in the Carpenter model (Equation (1)), the log-likelihood function cannot express analytically. It is evaluated numerically by solving the ordinary differential equations (ODEs) for a given parameter set and comparing the solution to observed data. As a result, the second derivatives of the log-likelihood cannot be obtained analytically. To overcome this, we use finite difference approximations, which provide a practical approach to estimate the required derivatives based only on numerical evaluations of the log-likelihood function.

Let *f* (***θ***) represent the log-likelihood function (Equation (4)), and let *δ* be a small perturbation. We approximate second partial derivatives using finite differences,

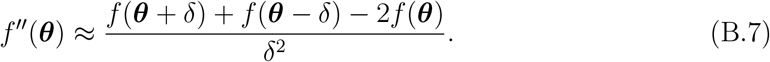

This provides estimates of the diagonal elements 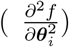 of the observed Fisher Information Matrix (Equation (B.6)), corresponding to the second derivative with respect to each individual parameter.

To estimate the off-diagonal elements, which correspond to the mixed second partial derivatives 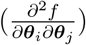 with respect to two parameters ***θ***_*i*_ and ***θ***_*j*_. The mixed second derivative is then approximated by

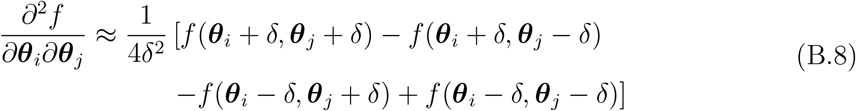

This expression is used to compute the off-diagonal elements of the observed FIM numerically.

## Appendix C. Additional Results

Here, we present additional results for different parameter sets that represent both non bistable and bistable regimes, capturing the oligotrophic and eutrophic states. These results are based on parameter estimates obtained via maximum likelihood estimation (Section 2.3), followed by identifiability analysis (Section 2.4) to assess parameter identifiability. We further examine the stability of each parameter set and determine the corresponding tipping points using the profile-wise analysis framework (Section 2.5).

### Appendix C.1. Regime Transition in Scenario 1

Scenario 1 represents the non bistable oligotrophic case (Figure 6), where the nonlinear parameters (*r, q*, and *p*_*h*_) are non-identifiable (Figure 8). Although the MLE is estimated (see Table 3), any combination of parameters within the confidence region can describe the system equally well. For example, adjusting the non-identifiable parameters to *r* = 1.2 g L^−1^ year^−1^, *q* = 7, and *p*_*h*_ = 0.8 g L^−1^, while keeping the identifiable parameters fixed, can result in a system that appears bistable. This case is represented as Scenario 4 (Figure C.1). Even though the true parameter set corresponds to a non bistable regime, the uncertainty in the nonlinear parameters can make the system appear as if it could cross into a bistable regime having alternative stable states at 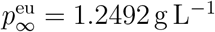. From a lake management perspective, if lake managers make decisions based only on the parameters estimated from the model (without considering the uncertainty or whether parameters are identifiable), they might misjudge how close the lake is to tipping point.

When we perform the profile likelihood analysis for the Scenario 4, the results indicate that the parameters *c, s*, and *σ* are identifiable (Figure C.1), within the narrow and well-defined peaks in their profile curves. In *c, s*, and *q* exhibit flat profile curves (Figure C.1), showing poor identifiability and large uncertainty. Stability analysis using the profile-wise framework further shows that, within the confidence region, certain parameter sets produce a non bistable system, where others lead to a bistable system. This mixture of stability behaviours implies that, it is not possible to determine whether the system is non bistable or bistable based on the available data and parameter estimates. These results highlight the importance of considering parameter uncertainty when interpreting the regime shifts, as non-identifiable parameters can lead to uncertain stability predictions.

**Figure C.1:**
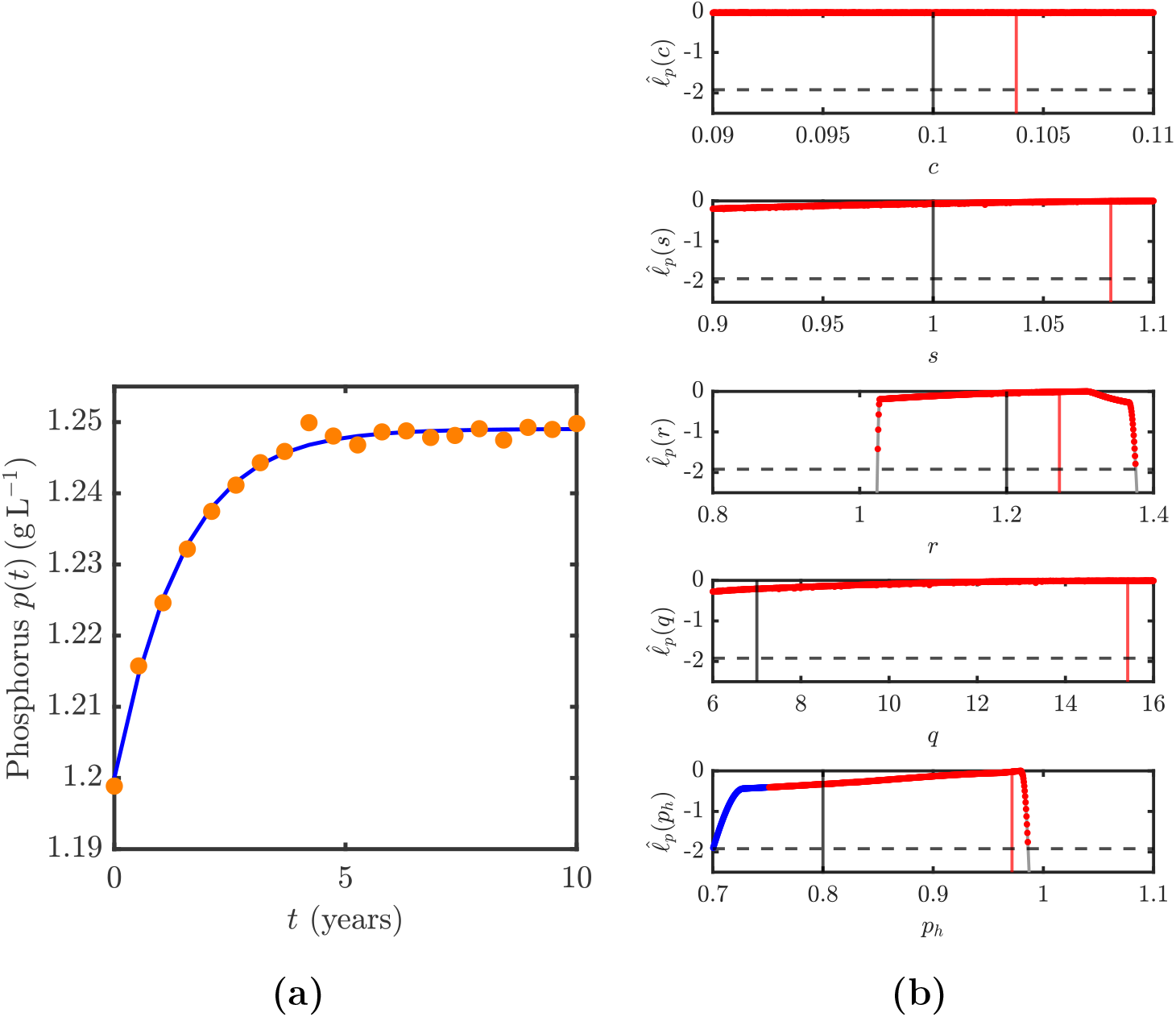
(a) Observed phosphorus concentrations (orange circles) representing the true behavior of the lake system along with the maximum likelihood estimate (MLE) trajectory (blue line). The true parameter values used to generate the synthetic dataset are: *c* = 0.1 g L^−1^, *s* = 1, *r* = 1.2, *q* = 7, *p*_*h*_ = 0.8 and *p*_0_ = 0.15. The MLE values for this scenario are *c* = 0.1038 g L^−1^, *s* = 1.0806, *r* = 1.2719, *q* = 15.4131, *p*_*h*_ = 0.9715 and *p*_0_ = 0.15. (b) Stability classification plots for Scenario 4 show the grid points where the normalized profile likelihood exceeds the threshold value of − 1.92 (dashed gray line). Blue circles indicate grid points classified as non bistable, while red circles indicate grid points classified as bistable.

#### Appendix C.2. Parameter Identifiability and Stability analysis for the eutrophic Scenarios

We focus on three primary oligotrophic Scenarios: one non bistable case and two bistable cases (Section 6) for the main calculations. However, in this section, we extend our framework to eutrophic Scenarios, applying the same methodology to three main eutrophic cases. As with the oligotrophic Scenarios, observations are simulated to reflect a long-term monitoring program, with biennial phosphorus measurements over 10 years (Guttal and Jayaprakash, 2007). For simplicity, we assume a highly precise monitoring program, setting the observational noise to *σ* = 0.001 g L^−1^.

The first Scenario in the eutrophic system, which consistently maintains a single stable upper eutrophic state near 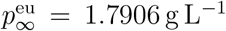 across varying initial phosphorus concentrations (Figure C.2 (a)), is represented as Scenario 5. This dataset is generated using model parameters *c* = 0.8 g L^−1^ year^−1^, *s* = 1 year^−1^, *r* = 1 g L^−1^ year^−1^, *q* = 8, and *p*_*h*_ = 1 g L^−1^, with the system initiated at *p*_0_ = 0.15 g L^−1^, following Guttal and Jayaprakash (2007). The second and third scenarios correspond to bistable systems within eutrophic regimes is known as Scenario 6 and Scenario 7, respectively. The Scenario 6, representing a bistable eutrophic regime (Figure C.2 (b)), the parameters are set to *c* = 0.5 g L^−1^ year^−1^, *s* = 1 year^−1^, *r* = 1 g L^−1^ year^−1^, *q* = 8, and *p*_*h*_ = 1 g L^−1^. The initial phosphorus concentration is chosen as *p*_0_ = 1.02 g L^−1^, stabilizing at the upper stable equilibria, again based on Guttal and Jayaprakash (2007). This regime features two stable equilibria, approximately 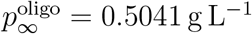 (lower stable equilibrium - oligotrophic) and 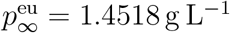, (upper stable equilibrium - eutrophic), separated by an unstable equilibrium near 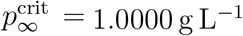. To demonstrate sensitivity to initial conditions, we also simulate the system with *p*_0_ = 1.02 g L^−1^, which is very close to the unstable equilibrium.

The Scenario 7, third Scenario of the eutrophic regime, represents a bistable eutrophic case (see Fig: C.2 (c)), but with perturbed parameters drawn randomly to simulate natural variability. The parameters are set to *c* = 0.1994 g L^−1^ year^−1^, *s* = 1.0648 year^−1^, *r* = 1.7337 g L^−1^ year^−1^, *q* = 5.9012, *p*_*h*_ = 0.8854 g L^−1^, with *p*_0_ = 1.02 g L^−1^. This setup also represents two stable equilibrium at approximately 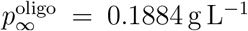, and 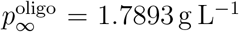. As the initial condition lies above but not too close to the unstable equilibrium 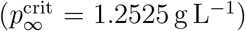, the trajectory again converges to the upper stable state. The qualitative difference between the three scenarios is that Scenarios 5 and 7 both look qualitatively similar, but Scenario 5 is generated from a non bistable system, and Scenario 7 being generated from a bistable system. In contrast, Scenario 6 is qualitatively distinct, with an initially polluted.

**Figure C.2:**
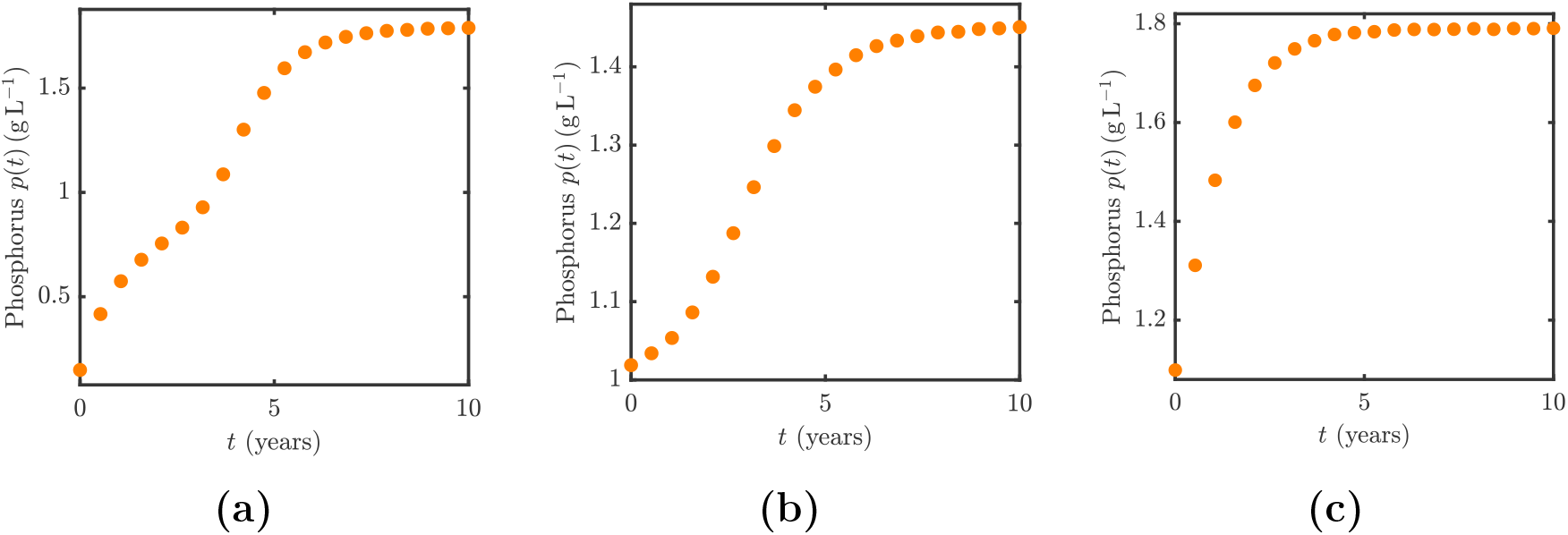
(a)–(c) Observed phosphorus data (orange circles), representing the actual phosphorus concentration behavior, are shown under eutrophic regimes and lower noise level (*σ* = 0.001 g L^−1^). Observations are recorded at discrete times *t*_0_ = 0, *t*_1_ = 0.5, …, *t*_20_ = 10 years. For the (a) Scenario 5 (non bistable–eutrophic regime), model parameters are *c* = 0.8 g L^−1^ year^−1^, *s* = 1 year^−1^, *r* = 1 g L^−1^ year^−1^, *q* = 8, *p*_*h*_ = 1 g L^−1^, with initial condition *p*_0_ = 0.15 g L^−1^. For the (b) Scenario 6 (bistable–eutrophic regime), parameters are *c* = 0.5 g L^−1^ year^−1^, *s* = 1 year^−1^, *r* = 1 g L^−1^ year^−1^, *q* = 8, *p*_*h*_ = 1 g L^−1^, with *p*_0_ = 1.02 g L^−1^. (c) Scenario 7 (Bistable–eutrophic regime) with perturbed parameters: *c* = 0.1994 g L^−1^ year^−1^, *s* = 1.0648 year^−1^, *r* = 1.7337 g L^−1^ year^−1^, *q* = 5.9012, *p*_*h*_ = 0.8854 g L^−1^, with *p*_0_ = 1.02 g L^−1^. Each plot converges to the corresponding upper stable states.

Following the approach used for the three main scenarios in the oligotrophic regime, we first determined the maximum likelihood estimates (MLEs) for each parameter in the eutrophic regime (Figure C.3). Table C.5 summarises both MLEs and corresponding 95% confidence intervals for these MLEs. We then conducted identifiability analysis (Section 2.4) to determine which parameters could be reliably estimated from the data.

**Figure C.3:**
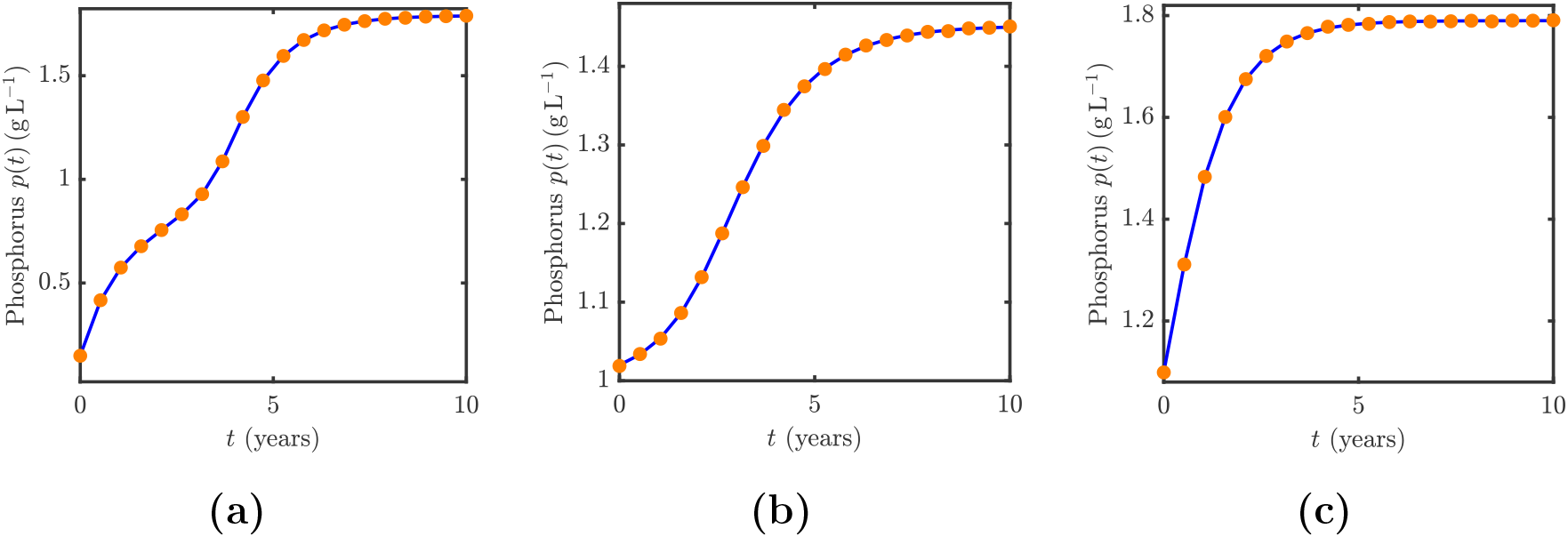
(a)–(c) Observed phosphorus concentrations (orange circles) are shown alongside the maximum likelihood estimate (MLE) trajectory (blue line), representing the expected mean phosphorus dynamics under non bistable and bistable regimes with a low observation noise level (*σ* = 0.001 g L^−1^). Observations are recorded at discrete time points: *t*_0_ = 0, *t*_1_ = 0.5, …, *t*_20_ = 10 years. For the (a) Scenario 5 (non bistable–eutrophic regime), model parameters are *c* = 0.8 g L^−1^ year^−1^, *s* = 1 year^−1^, *r* = 1 g L^−1^ year^−1^, *q* = 8, *p*_*h*_ = 1 g L^−1^, with initial condition *p*_0_ = 0.15 g L^−1^. For the (b) Scenario 6 (bistable–eutrophic regime), parameters are *c* = 0.5 g L^−1^ year^−1^, *s* = 1 year^−1^, *r* = 1 g L^−1^ year^−1^, *q* = 8, *p*_*h*_ = 1 g L^−1^, with *p*_0_ = 1.02 g L^−1^. (c) Scenario 7 (Bistable–eutrophic regime) with perturbed parameters: *c* = 0.1994 g L^−1^ year^−1^, *s* = 1.0648 year^−1^, *r* = 1.7337 g L^−1^ year^−1^, *q* = 5.9012, *p*_*h*_ = 0.8854 g L^−1^, with *p*_0_ = 1.02 g L^−1^.

**Table C.5:**
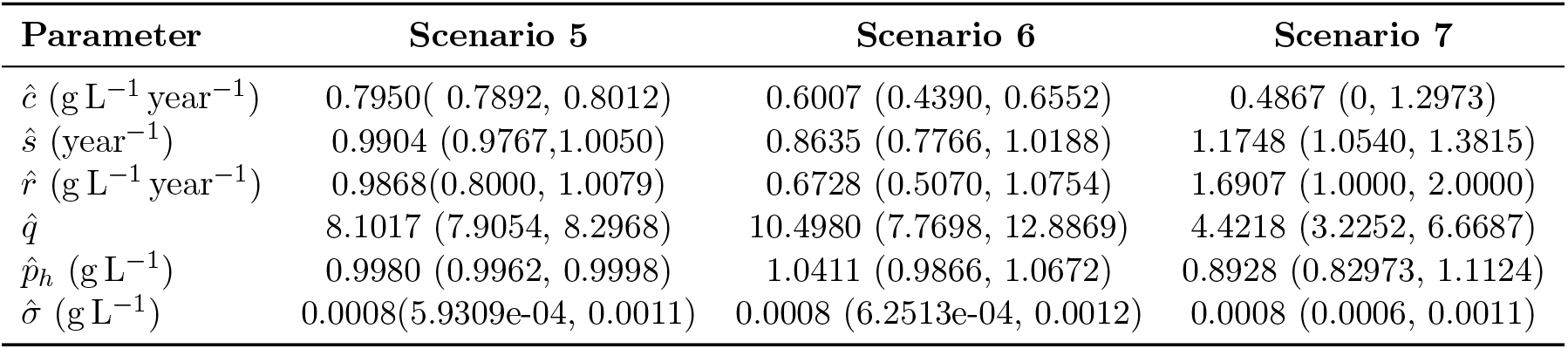
Maximum likelihood estimates (MLEs) and 95% Wald confidence intervals for the three eutrophic lake stability regimes.

| Parameter | Scenario 5 | Scenario 6 | Scenario 7 |
| --- | --- | --- | --- |
| $\hat{c} \text{ (g L}^{-1} \text{ year}^{-1}\text{)}$ | 0.7950( 0.7892, 0.8012) | 0.6007 (0.4390, 0.6552) | 0.4867 (0, 1.2973) |
| $\hat{s} \text{ (year}^{-1}\text{)}$ | 0.9904 (0.9767, 1.0050) | 0.8635 (0.7766, 1.0188) | 1.1748 (1.0540, 1.3815) |
| $\hat{r} \text{ (g L}^{-1} \text{ year}^{-1}\text{)}$ | 0.9868(0.8000, 1.0079) | 0.6728 (0.5070, 1.0754) | 1.6907 (1.0000, 2.0000) |
| $\hat{q}$ | 8.1017 (7.9054, 8.2968) | 10.4980 (7.7698, 12.8869) | 4.4218 (3.2252, 6.6687) |
| $\hat{p}_h \text{ (g L}^{-1}\text{)}$ | 0.9980 (0.9962, 0.9998) | 1.0411 (0.9866, 1.0672) | 0.8928 (0.82973, 1.1124) |
| $\hat{\sigma} \text{ (g L}^{-1}\text{)}$ | 0.0008(5.9309e-04, 0.0011) | 0.0008 (6.2513e-04, 0.0012) | 0.0008 (0.0006, 0.0011) |

Figures C.4, C.5, and C.6 present the univariate profile likelihood curves for three eutrophic Scenarios. Each profile includes a vertical line at the MLE and the 95% confidence interval derived from the profile likelihood analysis (Section 2.4). In Scenario 5 (non bistable eutrophic case; Figure C.3a), all model parameters are identifiable, as indicated by relatively narrow profile likelihood curves (Figure C.4). Scenario 6 (bistable case; Figure C.2b) also shows narrow curves for all parameters (Figure C.5), highlighting high identifiability. Here, the initial phosphorus concentration is close to the unstable equilibrium, which appears to improve parameter sensitivity. By contrast, in Scenario 7, the nonlinear recycling parameter *r* is non-identifiable, as indicated by a flat profile likelihood curve, while the remaining parameters remain identifiable within within uncertainty. In this case, the initial phosphorus concentration is not close to the unstable equilibrium, which may explain the reduced identifiability for *r*.

**Figure C.4:**
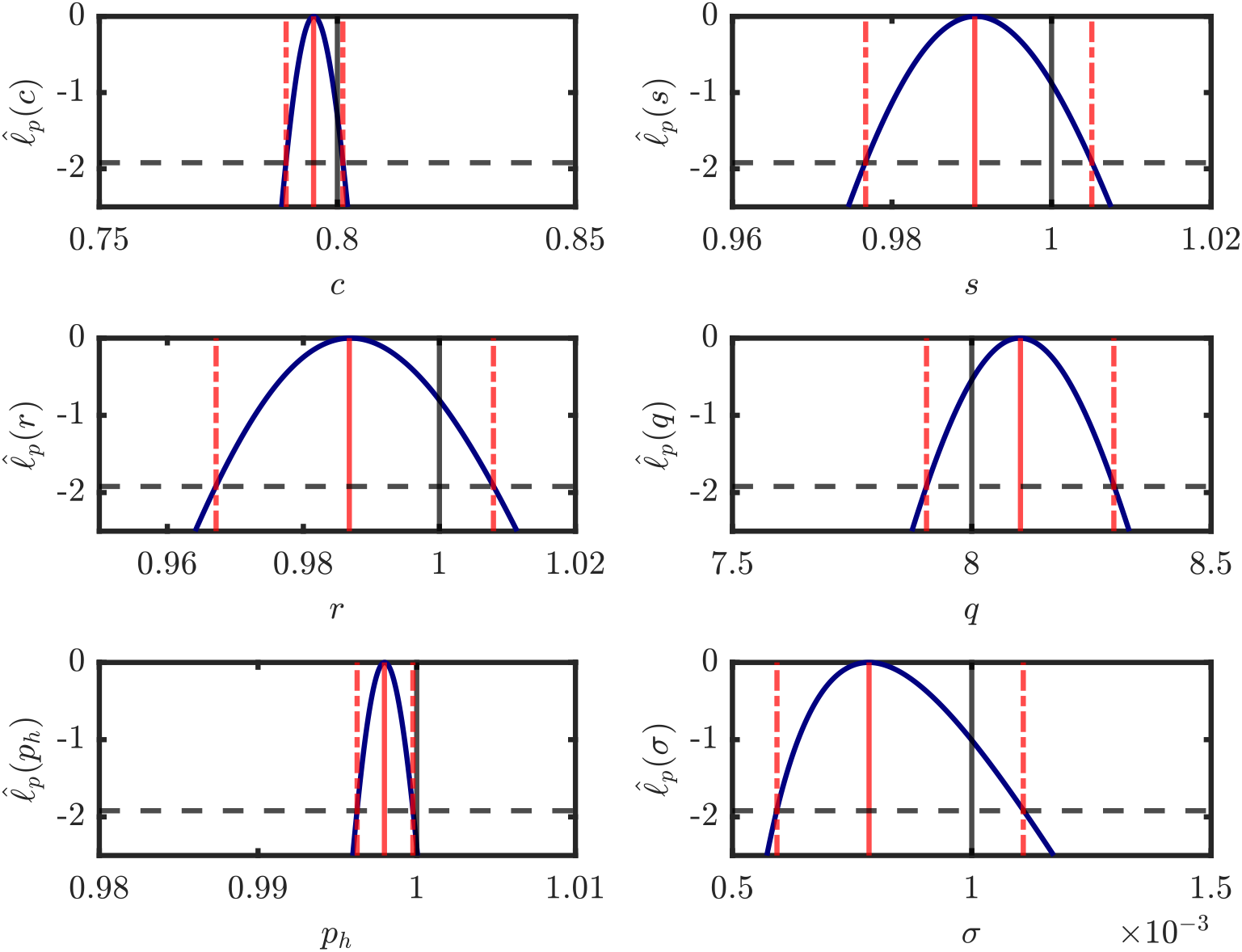
Profile likelihood plots for the Scenario 5 regime under lower noise levels (*σ* = 0.001 g L^−1^). Each plot shows the univariate profile likelihood curve (solid blue line), the MLE (red solid line), and the true parameter value (black solid line). The 95% confidence interval boundaries are shown (red dashed lines) based on the threshold of − 1.92 (black dashed line). The true parameter values used to generate the synthetic dataset are: *c* = 0.8 g L^−1^ year^−1^, *s* = 1 year^−1^, *r* = 1 g L^−1^ year^−1^, *q* = 8, *p*_*h*_ = 1 g L^−1^, with initial condition *p*_0_ = 0.15 g L^−1^.

**Figure C.5:**
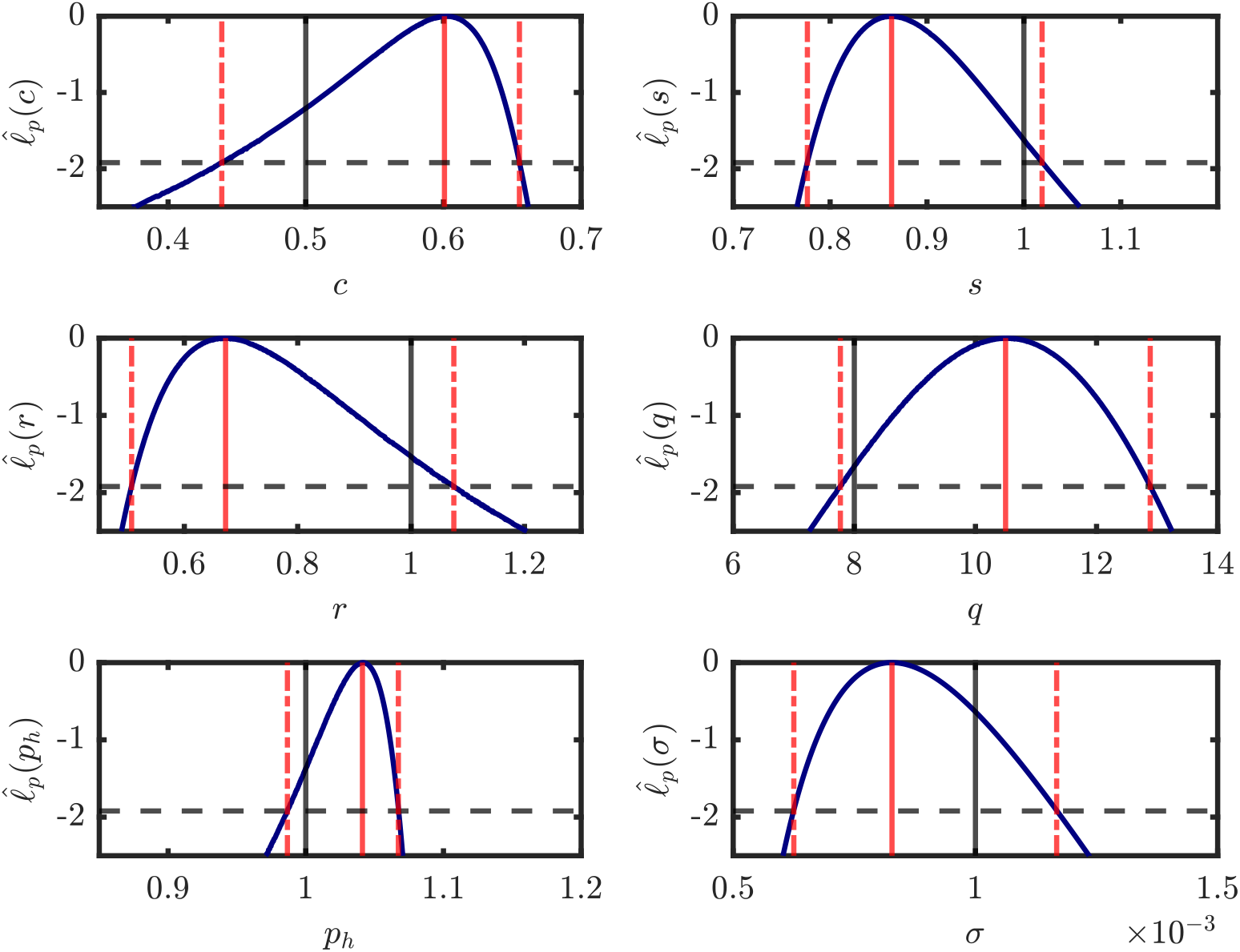
Profile likelihood plots for the Scenario 6 under lower noise levels (*σ* = 0.001 g L^−1^). Each plot shows the univariate profile likelihood curve (solid blue line), the MLE (red solid line), and the true parameter value (black solid line). The 95% confidence interval boundaries are shown (red dashed lines) based on the threshold of − 1.92 (black dashed line). The true parameter values used to generate the synthetic dataset are: *c* = 0.5 g L^−1^ year^−1^, *s* = 1 year^−1^, *r* = 1 g L^−1^ year^−1^, *q* = 8, *p*_*h*_ = 1 g L^−1^, with *p*_0_ = 1.02 g L^−1^.

**Figure C.6:**
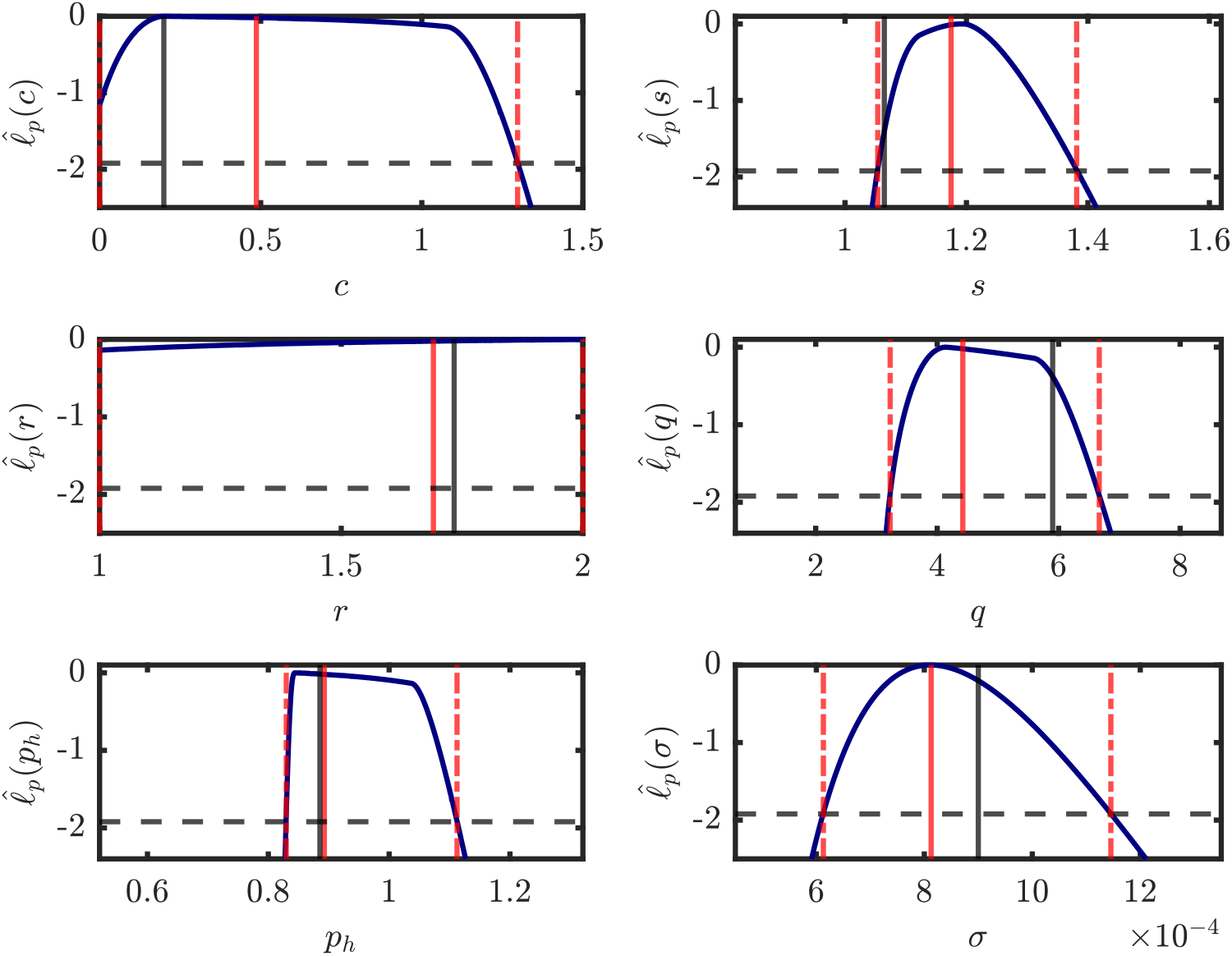
Profile likelihood plots for the Scenario 7 under lower noise levels (*σ* = 0.001 g L^−1^). Each plot shows the univariate profile likelihood curve (solid blue line), the MLE (red solid line), and the true parameter value (black solid line). The 95% confidence interval boundaries are shown (red dashed lines) based on the threshold of − 1.92 (black dashed line). The true parameter values used to generate the synthetic dataset are: *c* = 0.1994 g L^−1^ year^−1^, *s* = 1.0648 year^−1^, *r* = 1.7337 g L^−1^ year^−1^, *q* = 5.9012, *p*_*h*_ = 0.8854 g L^−1^, with *p*_0_ = 1.02 g L^−1^.

After completing the main calculation approach, we perform the identifiability analysis with a regime classification step. This is important, because it is challenging to determine whether a given scenario corresponds to a non bistable or bistable regime based solely on parameter estimates. To address this, we apply the statistical framework described in Section 2.5, which assesses the stability of each parameter set within the confidence intervals. The results for the three eutrophic scenarios are shown in Figure C.7.

The profile-wise analysis demonstrates that, in Scenario 5 (non bistable case), all parameter sets within the 95% confidence intervals lie within the non bistable region (Figure C.7a), consistent with the true system behaviour. In Scenario 6 (bistable case; Figure C.2b), where the initial condition is placed near the unstable equilibrium, the framework successfully identifies the system as bistable. The corresponding classification map (Figure C.7b) confirms that both the true parameter set and all parameter combinations within the confidence intervals fall within the bistable region (red circles). This suggests that closeness to the unstable equilibrium improves the detectability of bistability. In contrast, Scenario 7 (bistable case; Figure C.2c) presents greater uncertainty since initial condition is not close to the unstable equilibrium, and the profile-wise analysis reveals that parameter sets within the confidence intervals are split between non bistable and bistable classifications. As a result, even though the true system is bistable, the framework could not confirm bistability, highlighting the challenges of regime identification when the system operates far from the unstable equilibrium.

**Figure C.7:**
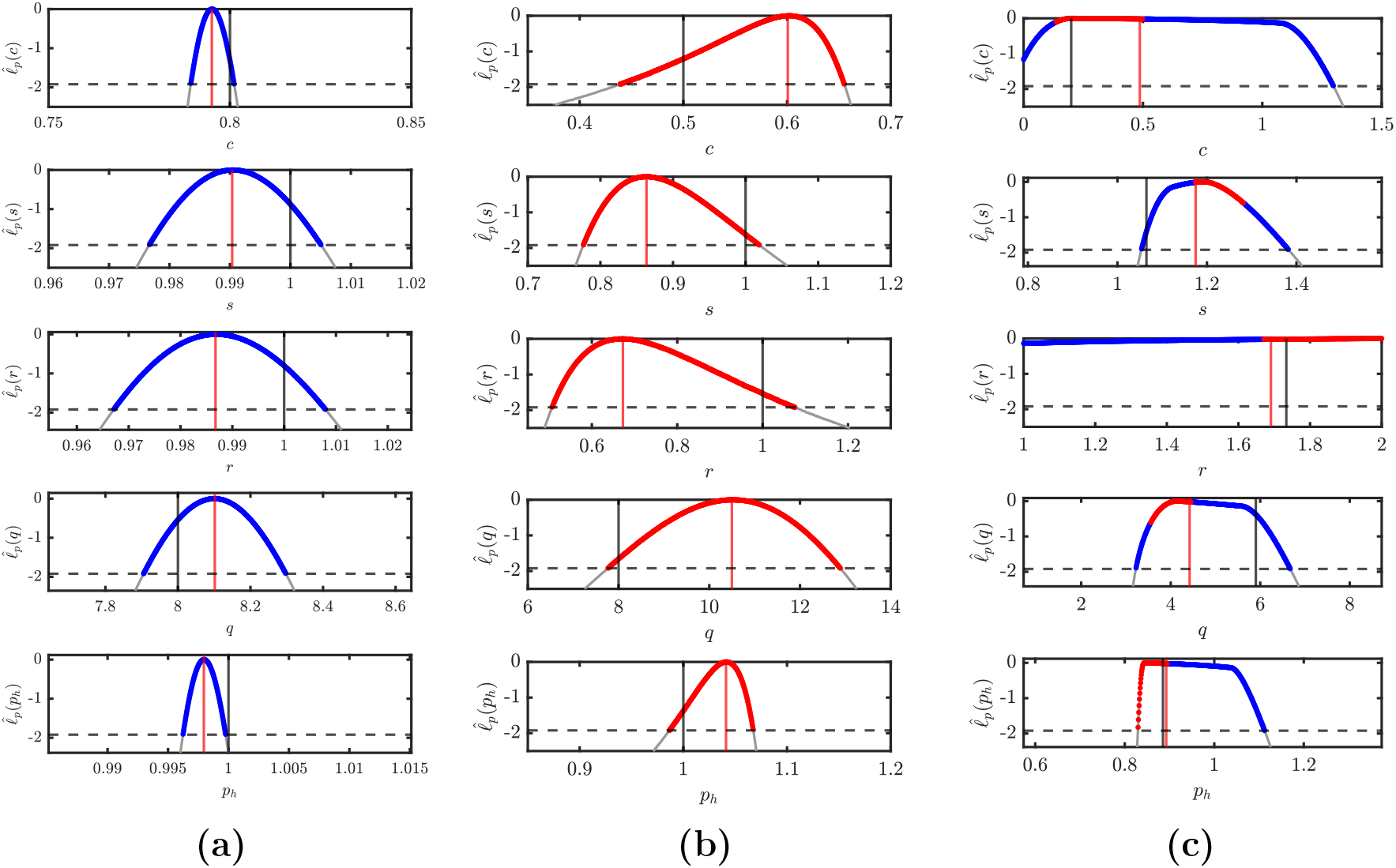
(a)–(c) shows stability classification plots for the three eutrophic scenarios. Each plot shows the grid points where the normalized profile likelihood exceeds the threshold value of − 1.92 (indicated by the dashed gray line). Blue circles indicate grid points classified as non bistable, while red circles indicate grid points classified as bistable. (a) Scenario 5 (non bistable–eutrophic regime), model parameters are *c* = 0.8 g L^−1^ year^−1^, *s* = 1 year^−1^, *r* = 1 g L^−1^ year^−1^, *q* = 8, *p*_*h*_ = 1 g L^−1^, with initial condition *p*_0_ = 0.15 g L^−1^. For the (b) Scenario 6 (bistable–eutrophic regime), parameters are *c* = 0.5 g L^−1^ year^−1^, *s* = 1 year^−1^, *r* = 1 g L^−1^ year^−1^, *q* = 8, *p*_*h*_ = 1 g L^−1^, with *p*_0_ = 1.02 g L^−1^. (c) Scenario 3 (Bistable–eutrophic regime) with perturbed parameters: *c* = 0.1994 g L^−1^ year^−1^, *s* = 1.0648 year^−1^, *r* = 1.7337 g L^−1^ year^−1^, *q* = 5.9012, *p*_*h*_ = 0.8854 g L^−1^, with *p*_0_ = 1.02 g L^−1^.

#### Appendix C.3. Parameter Identifiability and Stability analysis for the high noise level

In our study, we mainly focus on the lower noise level (*σ* = 0.001 g L^−1^), which represents a very precise monitoring program and is taken as the true noise level. At the same time, we also investigate a higher noise level (*σ* = 0.1 g L^−1^) to examine how stronger environmental variability influences both parameter identifiability and stability identifiability for both oligotrophic and eutrophic non bistable scenarios.

In this section, we focus on the non bistable oligotrophic scenario (Figure C.8 (a)) and the non bistable eutrophic scenario (Figure C.8 (a)) under the higher noise level (*σ* = 0.1 g L^−1^). For the oligotrophic case, synthetic data are generated using the parameter set *c* = 0.1 g L^−1^ year^−1^, *s* = 1 year^−1^, *r* = 1 g L^−1^ year^−1^, *q* = 8, *p*_*h*_ = 1 g L^−1^, with an initial condition *p*_0_ = 1.2 g L^−1^ (Figure C.8 (a)). For the eutrophic case, the parameter set is *c* = 0.8 g L^−1^ year^−1^, *s* = 1 year^−1^, *r* = 1 g L^−1^ year^−1^, *q* = 8, *p*_*h*_ = 1 g L^−1^, with an initial condition *p*_0_ = 0.15 g L^−1^ (Figure C.8 (b)).

Within the high noise, the observed phosphorus concentrations in both scenarios converge to their respective equilibrium values: 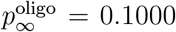 for the oligotrophic state and 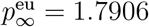 for the eutrophic state. However, the trajectories show much larger deviations around these equilibria (Figure C.8 (a)–(b)). These fluctuations occur due to the measurement noise obscures the true trend, causing the data to appear as the system derivates from equilibrium, even though the system is actually stable. As a result, the system may seem unstable or highly variable, which increases the difficulty of reliably estimating model parameters. When fitting the model using maximum likelihood estimation (MLE) (Section 2.3), the oligotrophic scenario determine parameter estimates as *c* = 0.0812 g L^−1^ year^−1^, *s* = 0.9098 year^−1^, *r* = 1.5000 g L^−1^ year^−1^, *q* = 16.0000, and *p*_*h*_ = 1.1629 g L^−1^ (Figure C.9 (a)). For the eutrophic scenario, the estimates are *c* = 0.7602 g L^−1^ year^−1^, *s* = 1.0671 year^−1^, *r* = 1.0931 g L^−1^ year^−1^, *q* = 8.5840, and *p*_*h*_ = 0.8867 g L^−1^ (Figure C.9 (b)). These values show the model’s best attempt to explain the noisy observations.

**Figure C.8:**
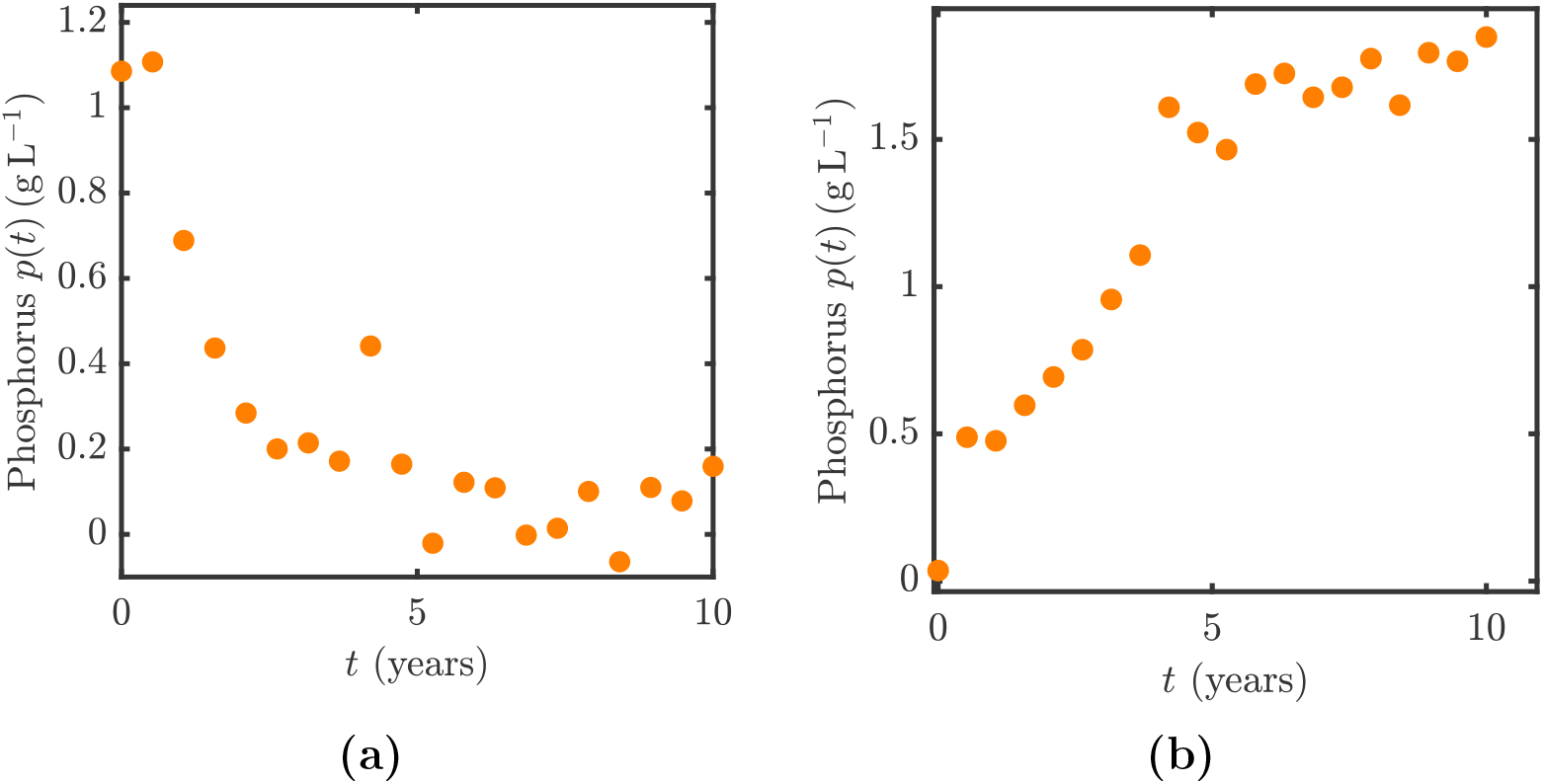
(a)–(b) Observed phosphorus data (orange circles) under the higher noise level (*σ* = 0.1 g L^−1^). Observations are recorded at discrete times *t*_0_ = 0, *t*_1_ = 0.5, …, *t*_20_ = 10 years. (a) Oligotrophic scenario generated with parameter values *c* = 0.1 g L^−1^ year^−1^, *s* = 1 year^−1^, *r* = 1 g L^−1^ year^−1^, *q* = 8, *p*_*h*_ = 1 g L^−1^, with initial condition *p*_0_ = 1.2 g L^−1^ which is stabilise at lower stable state. (b) Eutrophic scenario generated with parameter values *c* = 0.8 g L^−1^ year^−1^, *s* = 1 year^−1^, *r* = 1 g L^−1^ year^−1^, *q* = 8, *p*_*h*_ = 1 g L^−1^, with initial condition *p*_0_ = 0.15 g L^−1^ which is stablised at upper stable state.

**Figure C.9:**
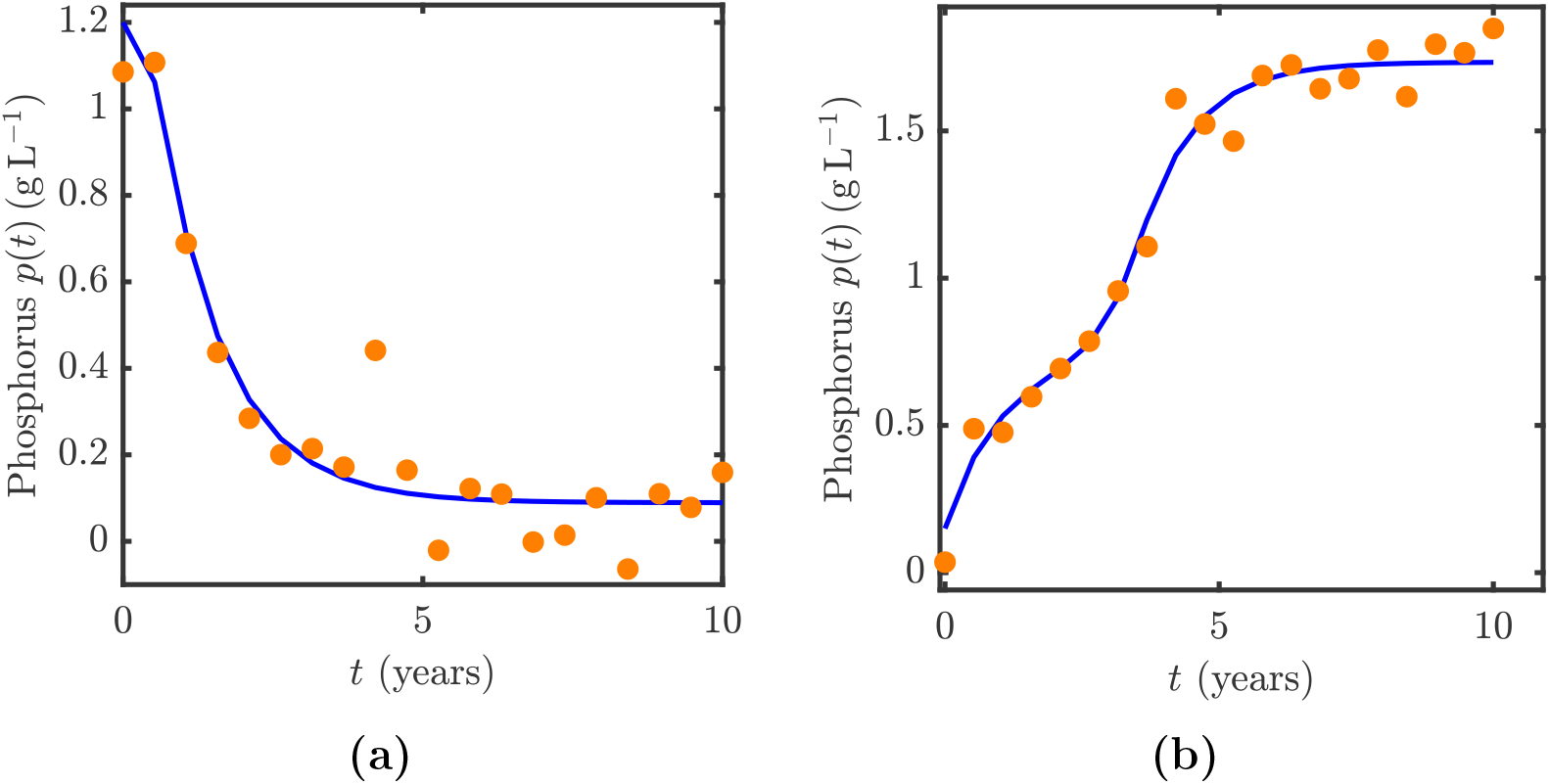
(a)–(b) Observed phosphorus data (orange circles) with the corresponding maximum likelihood estimation (MLE) trajectories (blue lines), representing the expected mean dynamics under the higher noise level (*σ* = 0.1 g L^−1^). Observations are recorded at discrete times *t*_0_ = 0, *t*_1_ = 0.5, …, *t*_20_ = 10 years. (a) Oligotrophic scenario: synthetic data are generated using the true parameter set *c* = 0.1 g L^−1^ year^−1^, *s* = 1 year^−1^, *r* = 1 g L^−1^ year^−1^, *q* = 8, *p*_*h*_ = 1 g L^−1^, with initial condition *p*_0_ = 1.2 g L^−1^. The MLE fit produce estimates of *c* = 0.0812 g L^−1^ year^−1^, *s* = 0.9098 year^−1^, *r* = 1.5000 g L^−1^ year^−1^, *q* = 16.0000, *p*_*h*_ = 1.1629 g L^−1^, with initial condition *p*_0_ = 1.2000 g L^−1^. (b) Eutrophic scenario: synthetic data are generated using the true parameter set *c* = 0.8 g L^−1^ year^−1^, *s* = 1 year^−1^, *r* = 1 g L^−1^ year^−1^, *q* = 8, *p*_*h*_ = 1 g L^−1^, with initial condition *p*_0_ = 0.15 g L^−1^. The MLE fit produce estimates of *c* = 0.7602 g L^−1^ year^−1^, *s* = 1.0671 year^−1^, *r* = 1.0931 g L^−1^ year^−1^, *q* = 8.5840, *p*_*h*_ = 0.8867 g L^−1^, with initial condition *p*_0_ = 0.1500 g L^−1^.

Once the model parameters are estimated, we perform identifiability analysis to assess whether they can be reliably determined from the data. Figures C.10 and C.11 show the univariate profile likelihood curves for the Carpenter model parameters (Equation (1)), with the maximum likelihood estimates superimposed. These results highlight that, although the parameters are identifiable under low noise conditions (*σ* = 0.001 g L^−1^) (Figure C.4), identifiability changes under higher noise levels (*σ* = 0.1 g L^−1^). For the oligotrophic scenario (Figure C.10), the profiles indicate that *c, s*, and *σ* remain identifiable, while *r, q*, and *p*_*h*_ are only one-sided identifiable. For the eutrophic scenario (Figure C.11), *c, s, p*_*h*_, and *σ* are identifiable, whereas the remaining parameters are only partially identifiable, indicating that they cannot be reliably estimated from the available data.

**Figure C.10:**
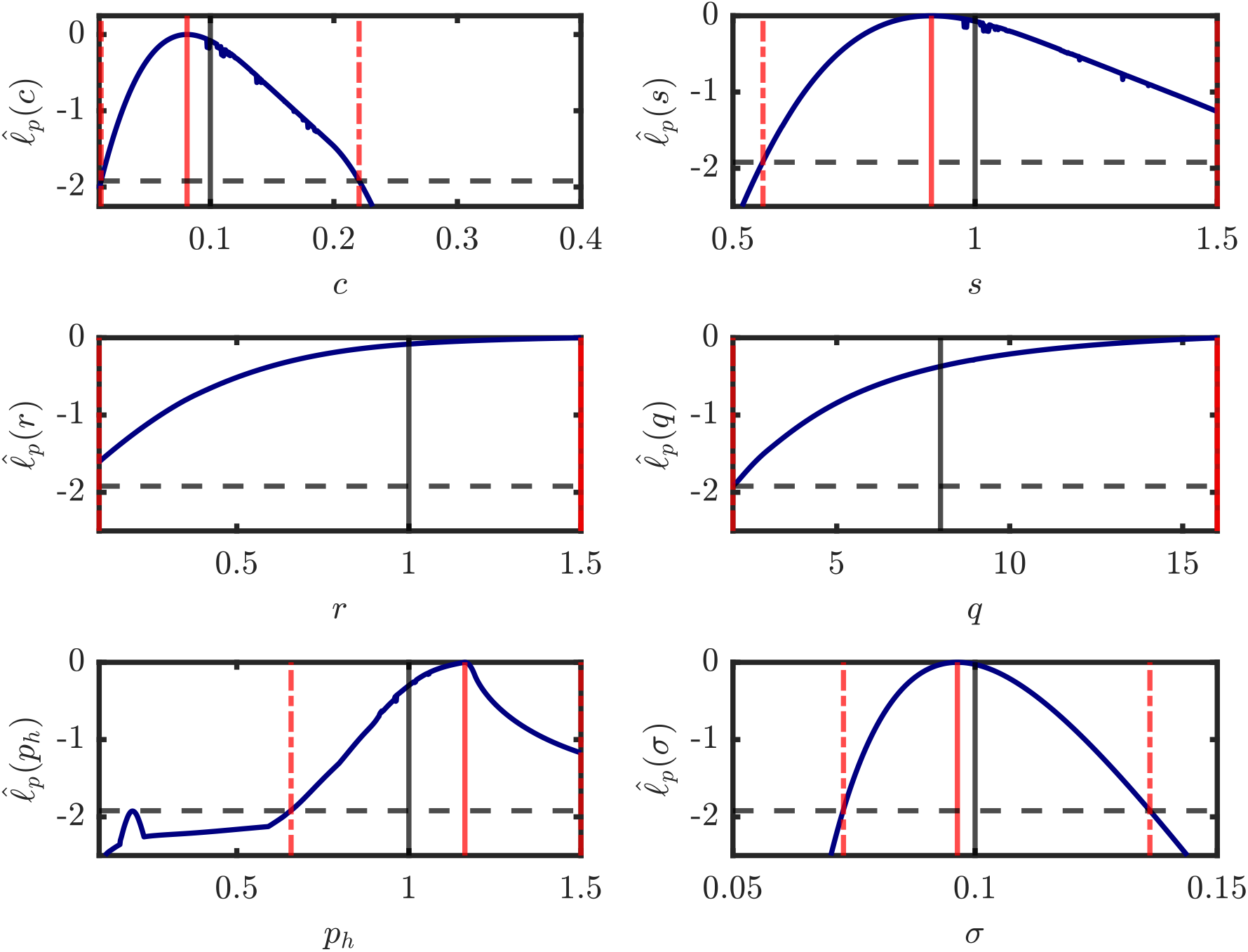
Profile likelihood plots for the higher noise levels (*σ* = 0.1 g L^−1^). Each plot shows the univariate profile likelihood curve (solid blue line), the MLE (red solid line), and the true parameter value (black solid line). The 95% confidence interval boundaries are shown (red dashed lines) based on the threshold of − 1.92 (black dashed line). The true parameter values used to generate the synthetic dataset for the oligotrophic are: *c* = 0.1 g L^−1^ year^−1^, *s* = 1 year^−1^, *r* = 1 g L^−1^ year^−1^, *q* = 8, *p*_*h*_ = 1 g L^−1^, with initial condition *p*_0_ = 1.2 g L^−1^.

**Figure C.11:**
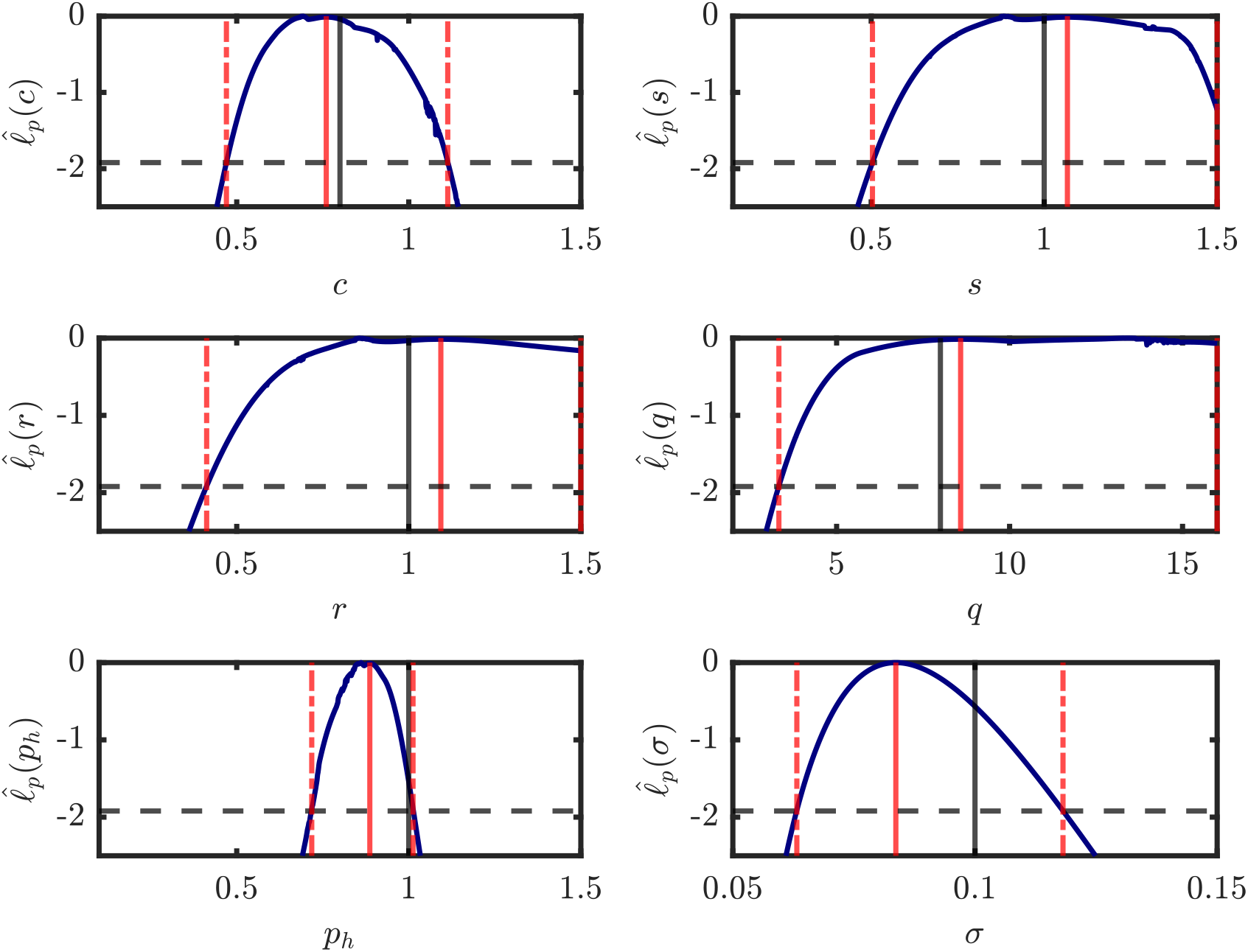
Profile likelihood plots for the higher noise levels (*σ* = 0.1 g L^−1^). Each plot shows the univariate profile likelihood curve (solid blue line), the MLE (red solid line), and the true parameter value (black solid line). The 95% confidence interval boundaries are shown (red dashed lines) based on the threshold of − 1.92 (black dashed line). Eutrophic scenario: synthetic data are generated using the true parameter set *c* = 0.8 g L^−1^ year^−1^, *s* = 1 year^−1^, *r* = 1 g L^−1^ year^−1^, *q* = 8, *p*_*h*_ = 1 g L^−1^, with initial condition *p*_0_ = 0.15 g L^−1^.

To further explore stability identification, we perform profile-wise analysis (Section 2.5) for both the oligotrophic and eutrophic scenarios (Figure C.8 (a)–(b)). As shown in Figure C.12 (a), the parameter sets within the 95% confidence interval for the oligotrophic scenario indicate mixed stability, exhibiting both non bistable and bistable behavior. Although the lower noise case (Section 11 (a)) clearly identifies the non bistable state as the ground truth, the higher noise level introduces greater uncertainty in stability classification. This increase in uncertainty occurs because measurement noise obscures the precise relationship between parameters and system behavior, making it harder to distinguish between parameter sets. In contrast, Figure C.12 (b) for the eutrophic scenario confirms the ground truth, consistent with the results shown in Figure 2.5. However, in the presence of high noise, the profile likelihoods for the parameters contains greater uncertainty, which limits the precision of both parameter estimation and the determination of system stability. Over-all, these results highlight that high observation noise can significantly affect confidence in stability predictions.

**Figure C.12:**
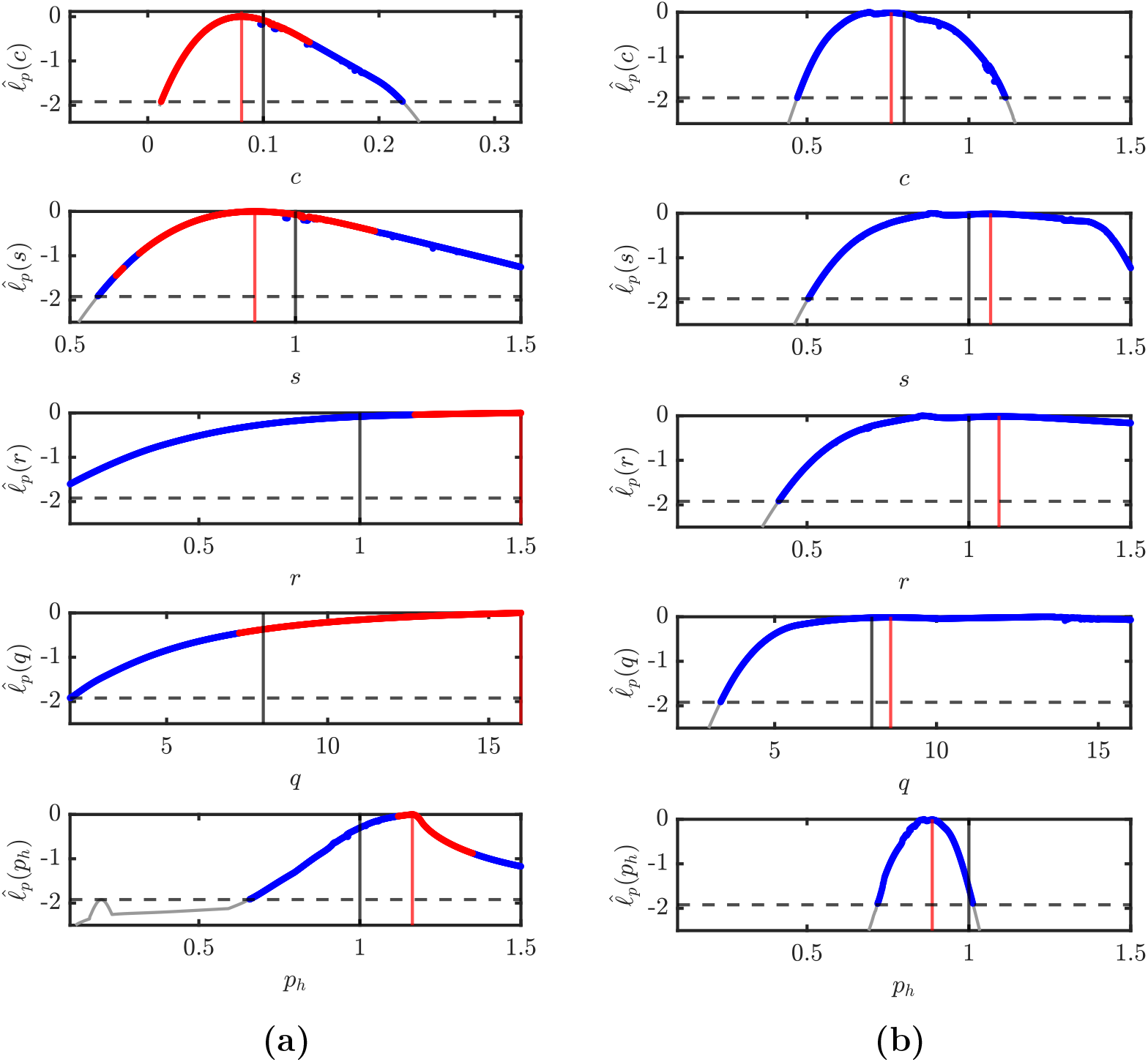
(a)–(b) show stability classification plots for the higher noise level (*σ* = 0.1 g L^−1^). Each plot shows the grid points where the normalized profile likelihood exceeds the threshold value of − 1.92 (dashed gray line). Blue circles indicate grid points classified as non bistable, while red circles indicate grid points classified as bistable. (a) Oligotrophic scenario: synthetic data are generated using the parameter set *c* = 0.1 g L^−1^ year^−1^, *s* = 1 year^−1^, *r* = 1 g L^−1^ year^−1^, *q* = 8, *p*_*h*_ = 1 g L^−1^. (b) Eutrophic scenario: synthetic data are generated using *c* = 0.8 g L^−1^ year^−1^, *s* = 1 year^−1^, *r* = 1 g L^−1^ year^−1^, *q* = 8, *p*_*h*_ = 1 g L^−1^, with initial condition *p*_0_ = 0.15 g L^−1^.

